# Comparative analysis of host-associated variation in *Phytophthora cactorum*

**DOI:** 10.1101/2021.02.22.432166

**Authors:** Charlotte F. Nellist, Andrew D. Armitage, Helen J. Bates, Maria K. Sobczyk, Matteo Luberti, Laura A. Lewis, Richard J. Harrison

**Author notes:** These authors contributed equally to the work (Charlotte F. Nellist, Andrew D. Armitage). MRC-IEU, Bristol Medical School, University of Bristol, Oakfield House, Oakfield Grove, Clifton, BS8 2BN, U.K. The Rhodes Trust, Rhodes House, Oxford, OX1 3RG, U.K.

## Abstract

*Phytophthora cactorum* is often described as a generalist pathogen, with isolates causing disease in a range of plant species. It is the causative agent of two diseases in the cultivated strawberry, crown rot (CR; causing whole plant collapse) and leather rot (LR; affecting the fruit). In the cultivated apple, *P. cactorum* causes girdling bark rots on the scion (collar rot) and rootstock (crown rot), as well as necrosis of the fine root system (root rot) and fruit rots. We investigated evidence for host specialisation within *P. cactorum* through comparative genomic analysis of 18 isolates. Whole genome phylogenetic analysis provided genomic support for discrete lineages within *P. cactorum*, with well supported non-recombing clades for strawberry CR and apple infecting isolates specialised to strawberry crowns and apple tissue. Isolates of strawberry CR are genetically similar globally, while there is more diversity in apple-infecting isolates. We sought to identify the genetic basis of host specialisation, demonstrating gain and loss of effector complements within the *P. cactorum* phylogeny, representing putative determinants of host boundaries. Transcriptomic analysis highlighted that those effectors found to be specific to a single host or expanded in the strawberry lineage are amongst those most highly expressed during infection of strawberry and give a wider insight into the key effectors active during strawberry infection. Many effectors that had homologs in other *Phytophthoras* that have been characterised as avirulence genes were present but not expressed in our tested isolate. Our results highlight several RxLR-containing effectors that warrant further investigation to determine whether they are indeed virulence factors and host-specificity determinants for strawberry and apple. Furthermore, additional work is required to determine whether these effectors are suitable targets to focus attention on for future resistance breeding efforts.

## INTRODUCTION

The *Phytophthora* genus serves as a model system for studying evolution of pathogenicity and resistance in plant pathosystems. Over 150 species have been named in the genus, with many pathogenic on plants (Yang et al., 2017). *Phytophthora* spp. are extremely effective plant pathogens that are able to disperse and infect hosts via the asexual, motile stage of their life cycle, zoospores, as well as able to resist and survive for many years in unfavourable conditions as thick-walled sexual oospores. Many *Phytophthora* spp. are specialised and only able to colonise one or a few host plants, for example *Phytophthora fragariae* which is thought to only colonise strawberry. Some species, such as *Phytophthora cactorum* (Lebert and Cohn) Schroet., are traditionally considered to be generalists and are able to cause disease on a broad range of plant species, including herbaceous and woody plants (Erwin and Ribeiro, 1996). Two examples of these hosts in the *Rosaceae* family, are the herbaceous cultivated strawberry (*Fragaria* x *ananassa*) and woody cultivated apple (*Malus* x *domestica*).

*P. cactorum* is the causative agent of two diseases in the cultivated strawberry, crown rot (CR; causing whole plant collapse; (Deutschmann, 1954) and leather rot (LR; affecting the fruit; (Rose, 1924). Both are considered major diseases of strawberry in temperate regions, with crop losses of up to 40% and 30% reported respectively for each disease (Ellis and Grove, 1983; Stensvand et al., 1999). The majority of commercial strawberries grown in the UK are grown under polytunnels or in glasshouses, on tabletops using soilless substrate, such as coir (coconut husk) (Boyer et al., 2016). *P. cactorum* is a particular problem in this production system due to ease of spread through the irrigation system via the motile asexual life-stage of *Phytophthora*, zoospores. Introduction into the irrigation system through asymptomatic infections in planting material represents the biggest risk to growers. Nursery propagation of strawberries is still based in the field, where the presence of resident pathogen inoculum in the soil along with latent infection in plants represent a severe threat to production. A study in 2018 of UK strawberry planting material (runners), commissioned by the Agriculture and Horticulture Development Board (AHDB), reported incidences of *P. cactorum* as high as 30% and great variation observed between batches of plants tested, but on average an incidence of 8-10% was recorded (Wedgwood et al., 2020).

In the cultivated apple, *P. cactorum* causes girdling bark rots on the scion (collar rot) and rootstock (crown rot), as well as necrosis of the fine root system (root rot) and fruit rots (Harris, 1991). *P. cactorum* affects nearly all apple growing regions of the world, debilitating the trees and leading to reduced fruit yield and eventual tree death (Alexander and Stewart, 2001). Due to the high costs associated with orchard establishment, losses due to *P. cactorum* can result in significant economic losses. As *P. cactorum* is homothallic (self-fertile), it produces sexual oospores which can survive for long periods in the soil, growing material and plant material, contributing to its importance as a worldwide threat to apple production.

*P. cactorum* diverged from other Clade 1 *Phytophthora* spp., *P. infestans* and *P. parasitica*, an estimated 221.4 Ma (138.6-342.4 million years ago; Yang et al., 2018) which is some 100 My (million years) earlier than the divergence of the Dryadoideae, Ammygdaloieae and Rosioideae and some 150-170 My before the emergence of the Fragariae (Zhang et al., 2017). *P. cactorum* specifically belongs to subclade 1a, along with *Phytophthora idaei* (Yang et al., 2017), a sister taxon and pathogen of another important *Rosaceae* crop, raspberry (*Rubus idaeus*). Despite such a broad host range being described for *P. cactorum*, host specialisation has been documented to particular plant species, including strawberry and apple (Seemüller and Schmidle, 1979). *P. cactorum* isolates originating from strawberry crowns were found to be less pathogenic on apple bark tissue than isolates originating from strawberry fruit or apple and vice versa, with apple and strawberry fruit isolates being less pathogenic on strawberry crowns. It was found that all pathotypes were able to cause disease in strawberry fruit (Seemüller and Schmidle, 1979). *P. cactorum* host specialisation has also been reported in other hosts, such as silver birch, peach and almond (Hantula et al., 1997, 2000; Lilja et al., 1998; Thomidis, 2003; Bhat et al., 2006). The specific genetic components responsible for host specialisation in *P. cactorum* are not known. Although, studies in both filamentous and bacterial pathogen systems support the model of effector repertoire being one of the key determinants in pathogen host range and nonhost resistance (Schulze-Lefert and Panstruga, 2011; Stam et al., 2014).

Genomic resources have recently become available for *P. cactorum*, with genomes released for isolates from beech, *Fagus sylvatica* (Grenville-Briggs et al., 2017), Chinese ginseng, *Panax notoginseng* (Yang et al., 2018) and strawberry, *F. x ananassa* (Armitage et al., 2018). *Phytophthora* spp. carry two major families of cytoplasmic effectors that are translocated into the host cell, RxLR and Crinklers, both have been characterised by specific motifs and consist of a rapidly evolving effector/modulating domain. The RxLR family of effectors are characterised by an N-terminal signal peptide, followed by an RxLR-s/dEER motif (Arginine, any amino-acid, Leucine, Arginine often followed by Serine/Aspartate, Glutamate, Glutamate, Arginine) and a variable C-terminal domain that may contain WY domain repeats (Wawra et al., 2012; Win et al., 2012). These effectors are renowned for suppressing host defence mechanisms through the manipulation of various aspects of plant defence (Anderson et al., 2015). The second family of cytoplasmic effectors are Crinklers (CRNs), named because of the leaf crinkling effect observed when expressed in host plants (Torto et al., 2003). They are characterised by an N-terminal LFLAK domain followed by a DWL domain, with a DI domain sometimes present between these two domains (Haas et al., 2009; Stam et al., 2013). These conserved domains are followed by variable C-terminal domains.

In addition to cytoplasmic effectors, *Phytophthora* spp. also deploy an arsenal of apoplastic effector proteins during infection, including a large number of hydrolytic enzymes, such as cutinases, glycoside hydrolases (GHs), pectinases, and proteases, which promote their infection and manipulation of the plant immune system (Armitage et al., 2018; Wang and Wang, 2018a, 2018b). They also encode members of extracellular phytotoxin families such as the conserved necrosis-inducing proteins (NLPs) and small cysteine-rich (SCR) proteins, for such as PcF (*Phytophthora cactorum* factor) (Orsomando et al., 2001) and INF1 (Kamoun et al., 1997).

Here, we further explore host specialisation and the basis of pathogenicity in *P. cactorum* by investigating multiple isolates collected from symptomatic strawberry crowns and fruit, as well as isolates from symptomatic apple tissue. We demonstrate that there are distinct lineages within *P. cactorum* showing adaptation to strawberry crowns and apple tissue. We show that these lineages are associated with unique effector complements and that these differential genes are highly expressed during plant infection. Taken together, this work elucidates key lineage specific effector genes playing roles in specialisation to strawberry and apple in *P. cactorum*.

## RESULTS

### HOST RANGE TESTING OF *P. CACTORUM*

#### *P. cactorum* isolates show specialisation to strawberry crowns and apple

A clear difference in pathogenicity on different plant tissues was observed between the isolates from the different *P. cactorum* pathotypes (**Figure 1** and **Supplementary Table 2**). All strawberry CR isolates were able to cause disease in strawberry crowns to varying degrees on the three different strawberry cultivars (**Figure 1A**). Of the two strawberry LR isolates, 11-40 was able to cause disease in both ‘Malling Opal’ and ‘Elsanta’, whereas 17-21 was only able to cause disease in the very susceptible ‘Malling Opal’ (**Figure 1A**). None of the apple isolates were able to cause disease in the strawberry crowns (**Figure 1A**). When a selection of *P. cactorum* isolates were screened on excised apple shoots, two of the three apple isolates were pathogenic, with R36/14 being the most pathogenic (**Figure 1B**). *P. cactorum* isolate 62471 was shown to be pathogenic on apple seedlings in a previous screen in 2018 (data not shown). Of the two strawberry LR isolates, 17-21 was more pathogenic on the apple shoots than 11-40 (**Figure 1B**). None of the strawberry CR isolates tested were able to cause disease in apple shoots (**Figure 1B**). The two strawberry LR isolates appear to have a broader host range than either strawberry CR or apple isolates and are able to cause disease in both strawberry crowns and apple shoots (**Figure 1**). All representative isolates from the three pathotypes of *P. cactorum* were able to colonise strawberry fruit (**Supplementary Table 2**). Isolate 62471 appeared to be the weakest apple isolate as it caused the lowest percentage infection in the strawberry fruit over the three experiments (**Supplementary Table 2**), coinciding with its weak pathogenicity on apple tissue. No disease symptoms were recorded in the strawberry tissue or apple tissue when challenged with the *P. idaei* isolates (**Figure 1** and **Supplementary Table 2**). Although, it should be noted that the *P. idaei* isolates were also tested on raspberry fruit but the results were inconclusive.

**FIGURE 1.**
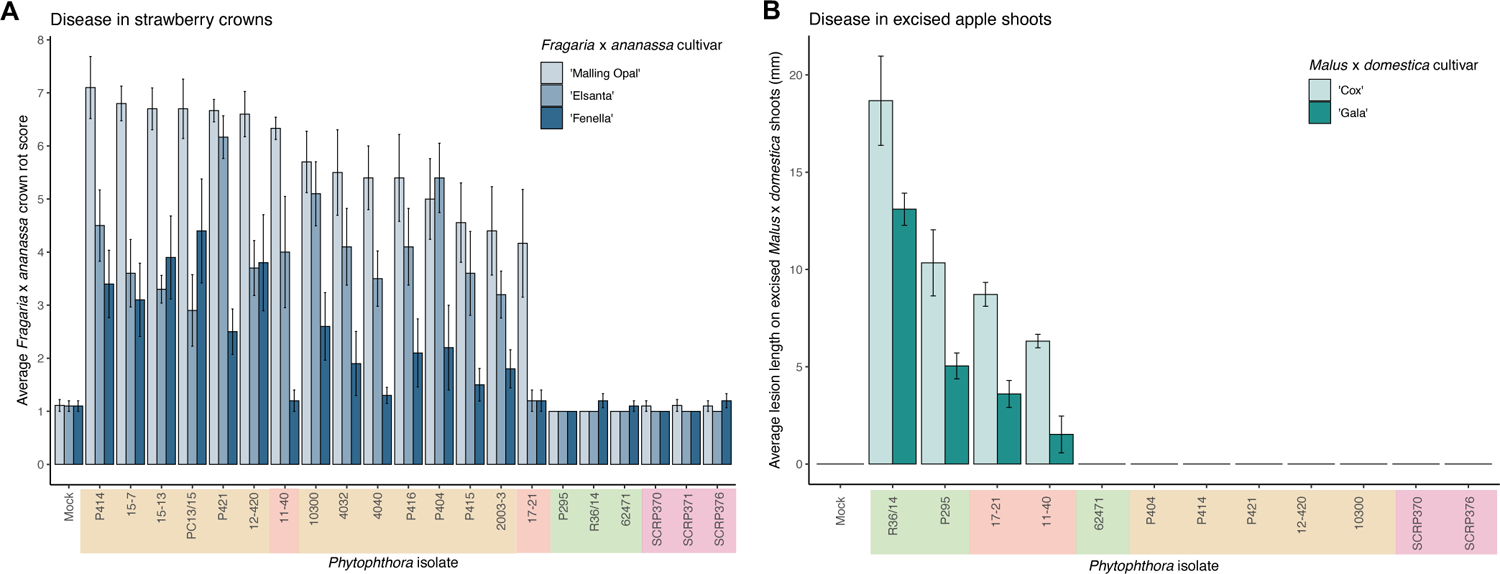
Phytophthora cactorum isolates show specialisation to strawberry crown and apple tissue. (A)-(B) Virulence of Phytophthora cactorum and Phytophthora idaei isolates on strawberry crowns and excised apple shoots by artificial inoculation of zoospores and mycelium, respectively. (A) Virulence of Phytophthora isolates on three Fragaria x ananassa cultivars, ‘Malling Opal’, ‘Elsanta’ and ‘Fenella’. Data are the mean of ten biological replicates ± se. (B) Virulence of Phytophthora isolates on two Malus x domestica cultivars, ‘Cox’ and ‘Gala’. Data are the mean of six biological replicates ± se.

#### An improved contiguous genome of *P. cactorum*

Here we present the best assembly to date of the plant pathogen *P. cactorum*. The SMRT data for the strawberry CR isolate P414 yielded an assembly of 66 Mb in 194 contigs. The other *P. cactorum* isolates, with the exception of P404, yielded *de novo* Illumina assemblies of 59.7-61.6 Mb in 4,452-6,726 contigs (**Table 1**), with isolate P404 larger and in a greater number of contigs, totalling 75.5 Mb in 20,136 contigs. The *P. idaei* genomes were a similar size to *P. cactorum* assemblies, 60.4-60.6 Mb in 4,720-5,356 contigs. Gene space between all assemblies was comparable, with 224-230 of 234 (95.7-98.3%) BUSCO genes both present and complete in the assemblies (**Table 1**), which were comparable to previous *Phytophthora* spp. sequencing projects, 91.5-94.4% for *P. cinnamomi* (Longmuir et al., 2018).

**TABLE 1.** Assembly statistics for 17 P. cactorum and three P. idaei isolates sequenced in this study. Isolate P414 represents results from long read sequencing technologies whereas other assemblies are a result of short read sequencing technology. Information on sequencing coverage, assembly size, percentage of bases repeatmasked are shown. Assembly completeness statistics, assessed using Alveolata-Stramenopiles Benchmarking Universal Single-copy Ortholog (BUSCO) genes, detail the number of complete (C) and total percentage of 234 core Eukaryotic genes present in assemblies.

| Features | *Phytophthora cactorum* |  |  |  |  |  |  |  |  |  |  | *Fragaria x ananassa*<br>fruit |  | *Malus x domestica* |  |  | *Phytophthora idaei*<br>*Rubus idaeus* |  |  |
| --- | --- | --- | --- | --- | --- | --- | --- | --- | --- | --- | --- | --- | --- | --- | --- | --- | --- | --- | --- |
|  | *Fragaria x ananassa*<br>crowns |  |  |  |  |  |  |  |  |  |  |  |  |  |  |  |  |  |  |
|  | P414 | 12-420 | 15-13 | 15-7 | 2003-3 | 4032 | 4040 | P404 | P415 | P421 | PC13/15 | 11-40 | 17-21 | 62471 | P295 | R36/14 | SCR370 | SCR371 | SCR376 |
| Coverage | 76.49 | 25.60 | 55.63 | 59.71 | 69.29 | 53.14 | 67.21 | 69.52 | 86.64 | 106.94 | 77.83 | 68.90 | 78.86 | 72.21 | 39.29 | 56.09 | 40.91 | 71.89 | 49.19 |
| No. of contigs | 194 | 5,509 | 5,421 | 5,457 | 5,467 | 6,726 | 5,326 | 2,0136 | 5,695 | 5,485 | 5,278 | 4,571 | 4,452 | 6,173 | 5,088 | 5,952 | 5,356 | 4,720 | 5,313 |
| Assembly size (Mb) | 66.0 | 59.9 | 60.7 | 60.8 | 60.8 | 61.6 | 60.7 | 75.5 | 60.8 | 60.5 | 60.0 | 61.1 | 59.7 | 59.8 | 61.3 | 60.7 | 60.5 | 61.7 | 60.6 |
| Longest contig (bp) | 1,749,730 | 272,591 | 358,283 | 300,516 | 353,051 | 299,428 | 238,735 | 346,862 | 313,236 | 251,913 | 346,832 | 288,536 | 347,645 | 285,881 | 357,485 | 345,260 | 247,223 | 295,942 | 230,545 |
| N50 (bp) | 645,375 | 37,498 | 41,248 | 43,376 | 40,273 | 38,202 | 39,295 | 36,598 | 40,742 | 34,103 | 39,779 | 43,828 | 49,722 | 36,408 | 48,651 | 36,465 | 39,576 | 46,381 | 39,406 |
| Masked (% bp) | 29.2 | 24.9 | 26.2 | 26.0 | 26.2 | 26.1 | 26.1 | 18.4 | 26.5 | 26.0 | 25.9 | 24.5 | 25.1 | 25.6 | 25.6 | 26.2 | 27.0 | 26.6 | 27.3 |
| BUSCO |  |  |  |  |  |  |  |  |  |  |  |  |  |  |  |  |  |  |  |
| C | 224 | 228 | 227 | 228 | 228 | 228 | 228 | 227 | 228 | 226 | 228 | 227 | 227 | 226 | 227 | 227 | 230 | 230 | 229 |
| % present | 95.7 | 97.4 | 97.0 | 97.4 | 97.4 | 97.4 | 97.4 | 97.0 | 97.4 | 96.6 | 97.4 | 97.0 | 97.0 | 96.6 | 97.0 | 97.0 | 98.3 | 98.3 | 97.9 |

#### Whole genome phylogeny supports resolution between *P. cactorum* pathotypes

A consensus phylogeny of 230 conserved single copy genes from the 22 *Phytophthora* isolates showed clear resolution between species, with *P. cactorum* and *P. idaei* isolates resolved into distinct clades (**Figure 2**). Resolution was also shown within *P. cactorum*, with the 13 strawberry CR isolates, including the previously sequenced CR isolate 10300, present in a distinct clade from the three apple isolates. The two strawberry LR isolates were placed into different clades. Strawberry LR isolate 11-40, which was more virulent on strawberry crowns, was placed in the same clade as the strawberry CR isolates. Whereas strawberry LR isolate 17-21, which was more virulent on apple, was placed in the same clade as the apple isolates. Interestingly, the publicly available *P. cactorum* isolate from *F. sylvatica* was genetically distinct from all other *P. cactorum* isolates (**Figure 2**). Within the strawberry clade, no evidence was observed for isolates being associated with geographical distribution, with isolates from the U.K., Netherlands, Norway and U.S.A., observed to be distributed throughout the clade.

**FIGURE 2.**
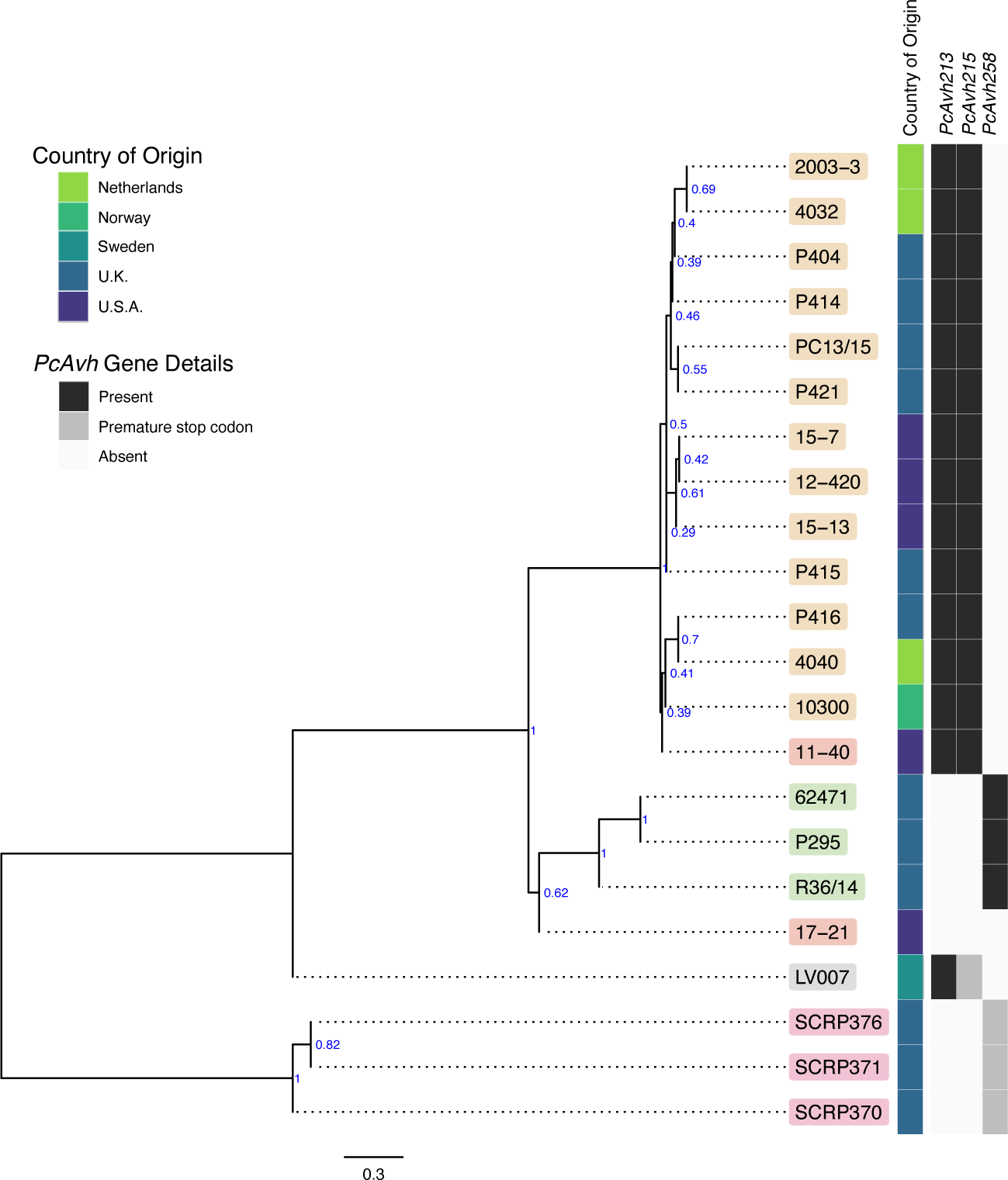
Apple and strawberry crown rot Phytophthora cactorum isolates form distinct clades. Maximum parsimony consensus phylogeny of 230 Alveolata-Stramenopiles Benchmarking Universal Single-Copy Ortholog (BUSCO) genes from 22 Phytophthora isolates. Branch support shown in blue. P. cactorum isolates shown within beige boxes and P. idaei isolates shown within grey boxes. Details of the host species isolated from, country of origin and presence/absence of a Phytophthora sojae Avh32 homolog and Phytophthora infestans Avr3a homolog are shown in the heatmap.

#### Population structure reflects predominant asexuality within diverging *P. cactorum* lineages

Population structure within *P. cactorum* was investigated using SNP variants predicted in relation to the strawberry CR isolate P414. *P. cactorum* SNP data was found to be best described by three non-recombining populations representing the strawberry lineage, apple lineage and the LR isolate 17-21 respectively (**Figure 3A**). Proportions of shared variants between these populations was found to be less than 0.1% in each isolate, indicating a lack of recombination between populations. Subpopulation structure also showed indications of asexuality when restricted to isolates within the strawberry lineage (**Figure 3B**). Isolates within the three subpopulations of the strawberry lineage showed high genetic identity to one another, with the number of heterozygous sites observed within a diploid individual being close to the genetic distance between two individuals from the same population (**Figure 3C**). For example, in the subpopulation consisting of 4040 and P416, 44 sites were found to be heterozygous within isolate 4040, whereas the total number of SNPs that showed variability between isolate 4040 and P416 (including heterozygous and homozygous sites) was observed to be 79 (**Figure 3C**). This was in contrast to the number of SNPs differing between isolates between subpopulations, where isolate 4040 differed by a total of 1,054 SNPs to its next closest isolate, 15-7, from another subpopulation (**Figure 3C**).

**FIGURE 3.**
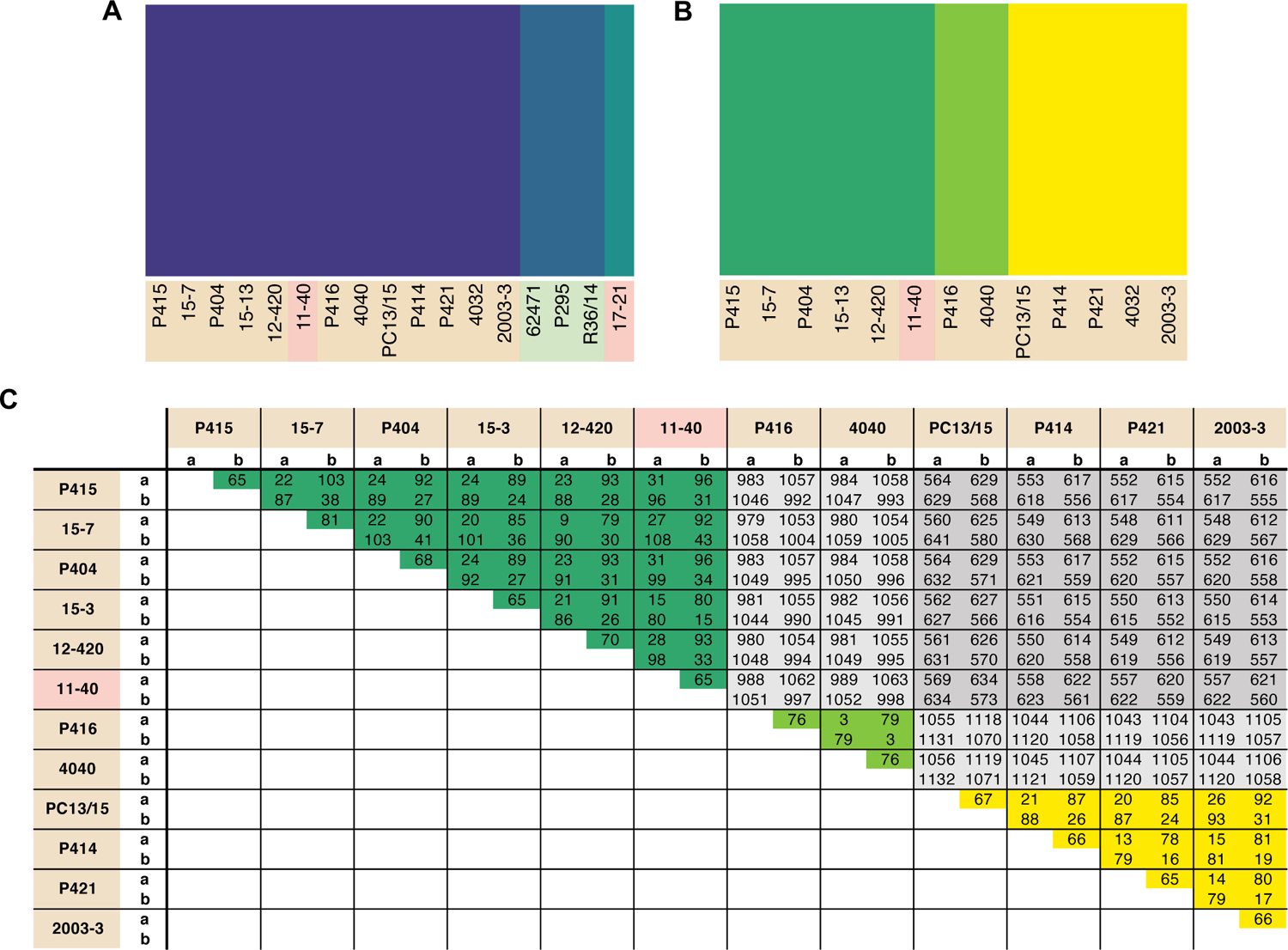
Phytophthora cactorum isolates split into three populations with predominant asexuality. Population analysis of high quality, biallelic SNP sites split the P. cactorum isolates into three populations, admixture of variants was not observed within P. cactorum, or within a single lineage of P. cactorum indicating predominant asexuality. DISTRUCT plot of fastSTRUCTURE (Raj et al., 2014) results carried out on (A) all sequenced isolates of P. cactorum and (A) strawberry lineage isolates. Data was best described by three “populations” in both datasets. (C) SNP calling was performed in relation to reference isolate P414 for the P. cactorum strawberry lineage. The number of differing sites are shown between the two haplotypes from each isolate; a, representing only homozygous variants and b, representing both homozygous and heterozygous variants in relation to the P414 reference genome.

### IDENTIFICATION OF POPULATION-, PATHOTYPE- AND SPECIES-LEVEL VARIATION IN *P. CACTORUM* EFFECTORS

#### *P. cactorum* possesses an expanded effector repertoire in comparison to *P. idaei*

The predicted proteome of *P. cactorum* CR isolate P414 totalled 29,913 proteins encoded by 29,552 genes. Additional isolates were predicted to carry a similar number of proteins 25,449-29,955 and genes 24,856-29,124, with the exception of P404 (**Table 2**). Within *P. cactorum*, strawberry and apple isolates had similar numbers of predicted genes and effectors. However, *P. idaei* isolates carried a reduced predicted effector repertoire (**Table 2**), with fewer secreted carbohydrate active enzymes (CAZYmes), CRNs, Elicitins, necrosis-inducing proteins (NLPs), glucanase inhibitors and kazal protease inhibitors predicted than *P. cactorum* isolates.

**TABLE 2.**
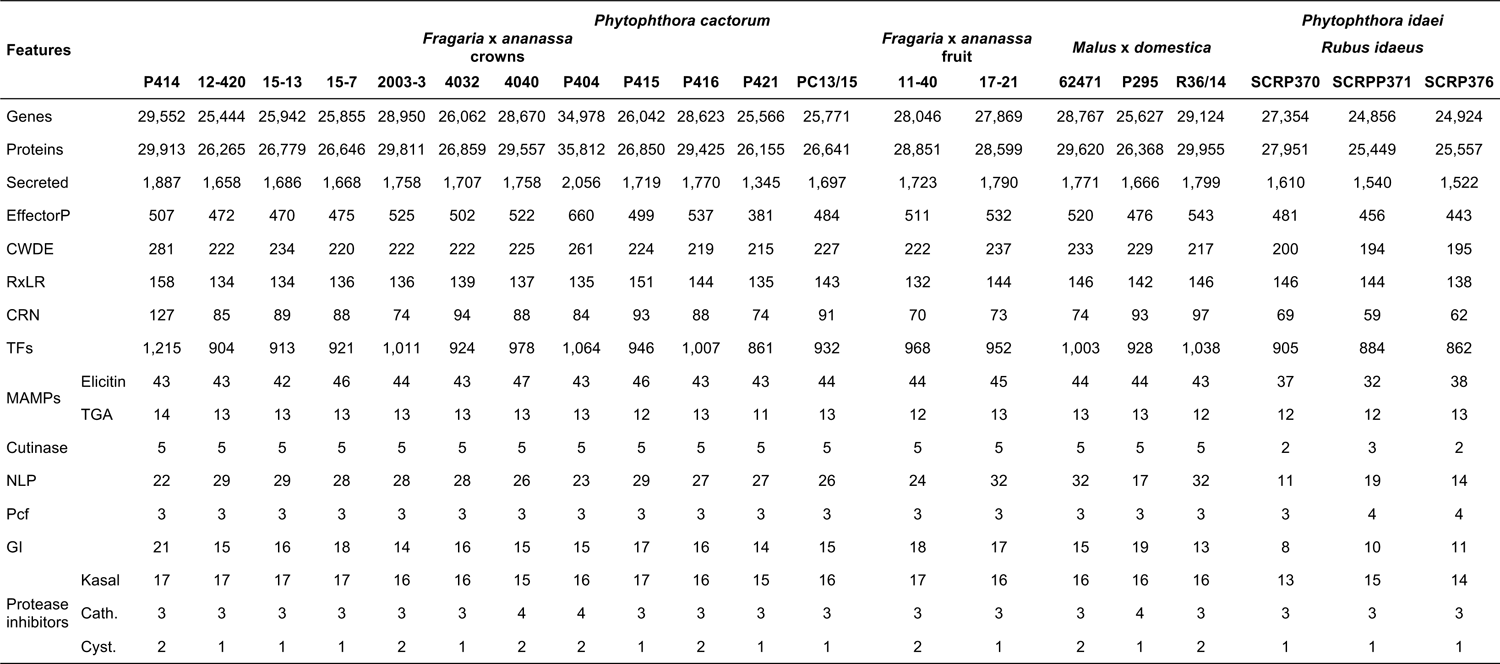
Predicted gene models and effectors. Total number of gene models and the predicted proteins that they encode are shown. Numbers of secreted proteins, secreted cell wall degrading enzymes (CWDE), RxLR and crinkler (CRN) family cytoplasmic effectors are shown, along with predicted transcription factors (TFs). Host defence triggering (MAMP) family sterol binding (Elicitn) and transglutanimase (TGA) proteins are shown as well as apoplastic effector families including necrosis inducing proteins (NLP), Phytophthora cactorum factor (PcF), and protease inhibitors (kazal-, cathepsin- and cystatin-acting).

#### Orthology analysis identifies gene expansion and contraction associated with phylogenetic lineage

The total set of 157,038 predicted proteins from the 20 sequenced isolates, as well as the proteome of *P. cactorum* isolate 10300 from (Armitage et al., 2018), were clustered into 22,572 orthogroups. Orthogroups showing a consistent pattern of expansion/contraction by phylogenetic clade were identified (**Figure 4**). This allowed investigation into expansion/contraction events associated with the strawberry CR and apple lineages of *P. cactorum*.

**FIGURE 4.**
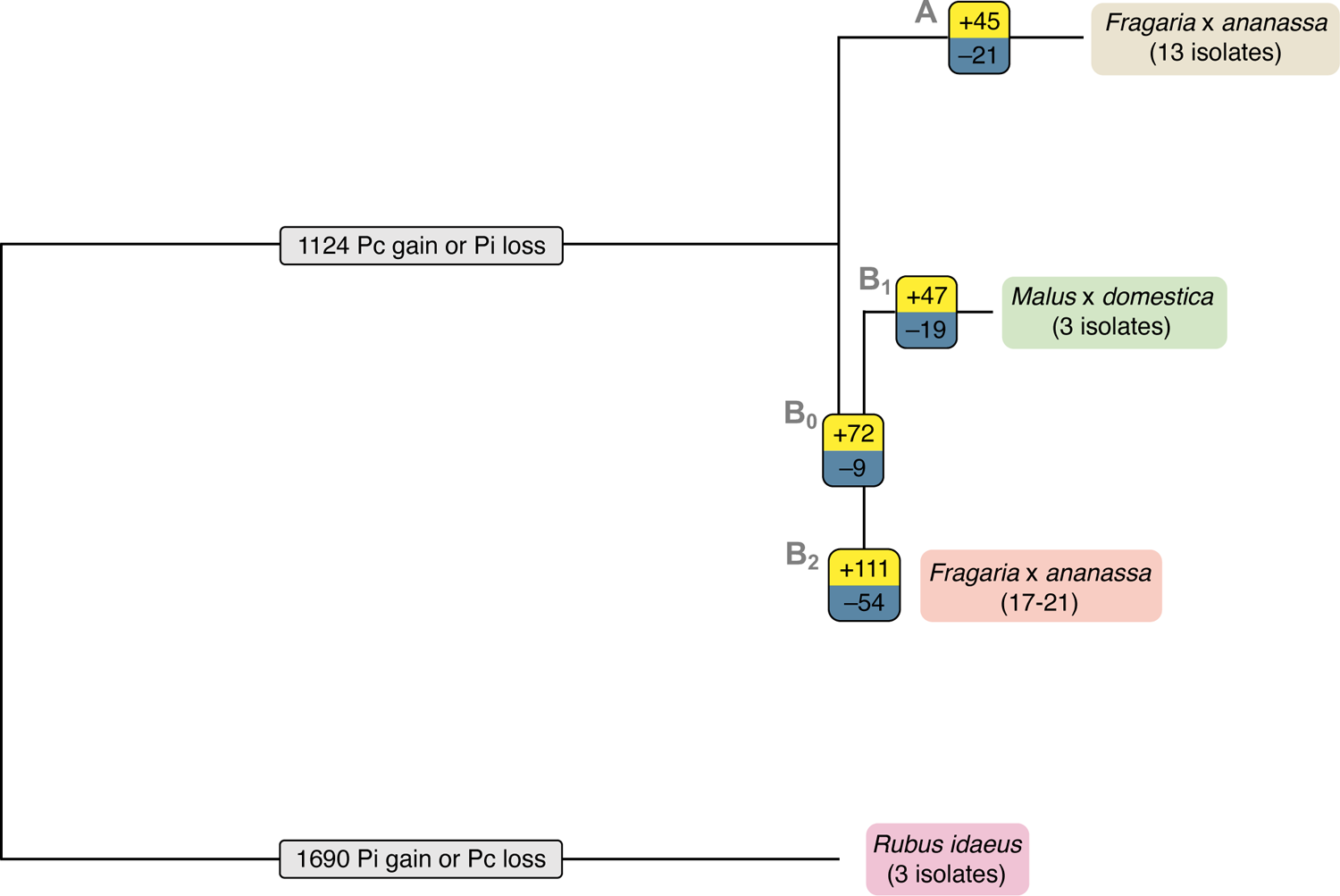
Expansion and contraction of orthogroups indicates possible roles in Phytophthora cactorum host specialisation. Expansion and contraction of orthogroups plotted onto simple phylogeny. The number of expanded orthogroups of each branch are shown in yellow, the number of contracted orthogroups of each branch are shown in blue and the numbers of orthogroups with unsure direction of expansion or contraction are shown in light grey. The strawberry crown rot lineage (branch A) includes isolates P414, P404, P415, P416, P421, 4032, 4040, 12-420, 2003-3, 10300, 15-7, 15-13 and PC13/15. The apple lineage (branch B1) includes isolates R36/14, P295 and 62471.

The 45 orthogroups expanded in the strawberry CR lineage (Branch A; **Figure 4**), represented a total of 65 genes from P414 (**Supplementary Table 3**). This included a number of potential effector candidates, notably two RxLRs (*Pcac1_g24384* and *Pcac1_g22827*) and an additional secreted protein (*Pcac1_g6287*). BLAST searches confirmed the absence of *Pcac1_g24384* and *Pcac1_g22827* in the apple lineage to be the result of these regions being absent from assemblies rather than genes not being predicted in those genomes. The strawberry CR lineage was found to have contracted across 21 orthogroups (Branch A; **Figure 4**), representing 33 genes in P414 (**Supplementary Table 3**). This included three RxLRs (*PC123_g16852*, *PC123_g26877* and *PC123_g27632*) and two additional secreted proteins (*PC123_g10425* and *PC123_g25979*) with effector-like structure that were lost in relation to the wider phylogeny.

The apple isolates and 17-21 lineage harboured greater diversity, with 119 orthogroups expanded (Branches B_0_, B_1_; **Figure 4**), representing a total of 241 genes (**Supplementary Table 3**). This included five RxLR (*PC123_g15654*, *PC123_g17462*, *PC123_g19522*, *PC123_g24792* and *PC123_g25079*) and two CRN (*PC123_g21108* and *PC123_g24736*) candidates, as well as seven secreted proteins (*PC123_g10425*, *PC123_g10510*, *PC123_g11377*, *PC123_g14333*, *PC123_g24080*, *PC123_g27245* and *PC123_g28487*), six of which had an effector-like structure (**Supplementary Table 3**). BLAST searches confirmed the absence of these genes in the strawberry CR lineage to be a result of these regions being absent from assemblies rather than genes not being predicted in those genomes. A single RxLR (*PC123_g19522*) and CRN (*PC123_g24736*) candidate were identified unique to apple isolates (Branch B_1_; **Figure 4**). *PC123_g19522* (hereafter *PcAvh258*; all RxLR homologs summarised in **Supplementary Table 4**) was found to have 58% pairwise aa homology (downstream of the signal peptide, aa 24-139) to *P. infestans Avr3a* (GenBank AEH27535.1; aa 22-147) (Armstrong et al., 2005). Further investigation of *PC123_g24792* noted a difference between the apple isolate homologues and that in 17-21. The homologue in the three apple isolates was truncated (69 aa; hereafter *PcAch246t*) compared to *PC128_g25726* in 17-21 (195 aa; hereafter *PcAvh246*). A non-synonymous SNP introduced at G210A resulted in a stop codon.

Contraction of 28 orthogroups was observed in the apple lineage (Branches B_0_, B_1_; **Figure 4**), representing 45 genes. This included only one RxLR candidate (*Pcac1_g13631*) and three additional secreted proteins (*Pcac1_g3068*, *Pcac1_g3069* and *Pcac1_g25117*), indicating a substantial increase in the effector complement of this lineage.

Overall, these results show that the apple lineage within *P. cactorum*, harbours greater diversity in effector complement than the strawberry CR lineage. RxLRs and CRNs were represented in the expanded and contracted gene families, as well as other unannotated proteins with an effector-like structure. However, other commonly observed *Phytophthora* pathogenicity factors such as secreted CAZYmes, elicitins and protease inhibitors were notably absent from these groups.

#### Polarising of SNP and indel variants identifies putative host specialisation events in effectors

Further variants were determined through identification of SNP, indel and small structural variants (insertions and duplications) from all sequenced isolates in comparison to strawberry CR isolate P414 (**Table 3**). Polarising of non-synonymous SNPs and indels to the outgroup *P. idaei* allowed identification of those variants that differed at the species level (private to *P. cactorum*), at the pathotype level (private to apple or strawberry CR isolates), or at the population level. Those variants at the pathotype level were investigated to identify potential signatures of host adaptation. In total, variants were observed in 21 RxLR and 12 CRN genes at the pathotype level (**Supplementary Table 5**). Of the RxLRs, eight genes contained non-synonymous variants unique to strawberry isolates and 13 unique to apple isolates. Of the CRNs, six genes contained non-synonymous variants unique to strawberry isolates and eight unique to apple isolates (with two genes containing unique variants in both), which in addition to the gene family expansion/contractions described above potentially represent host adaptation events or determinants of host boundaries between pathotypes, or simply functionally neutral mutations fixed due to drift.

**TABLE 3.**
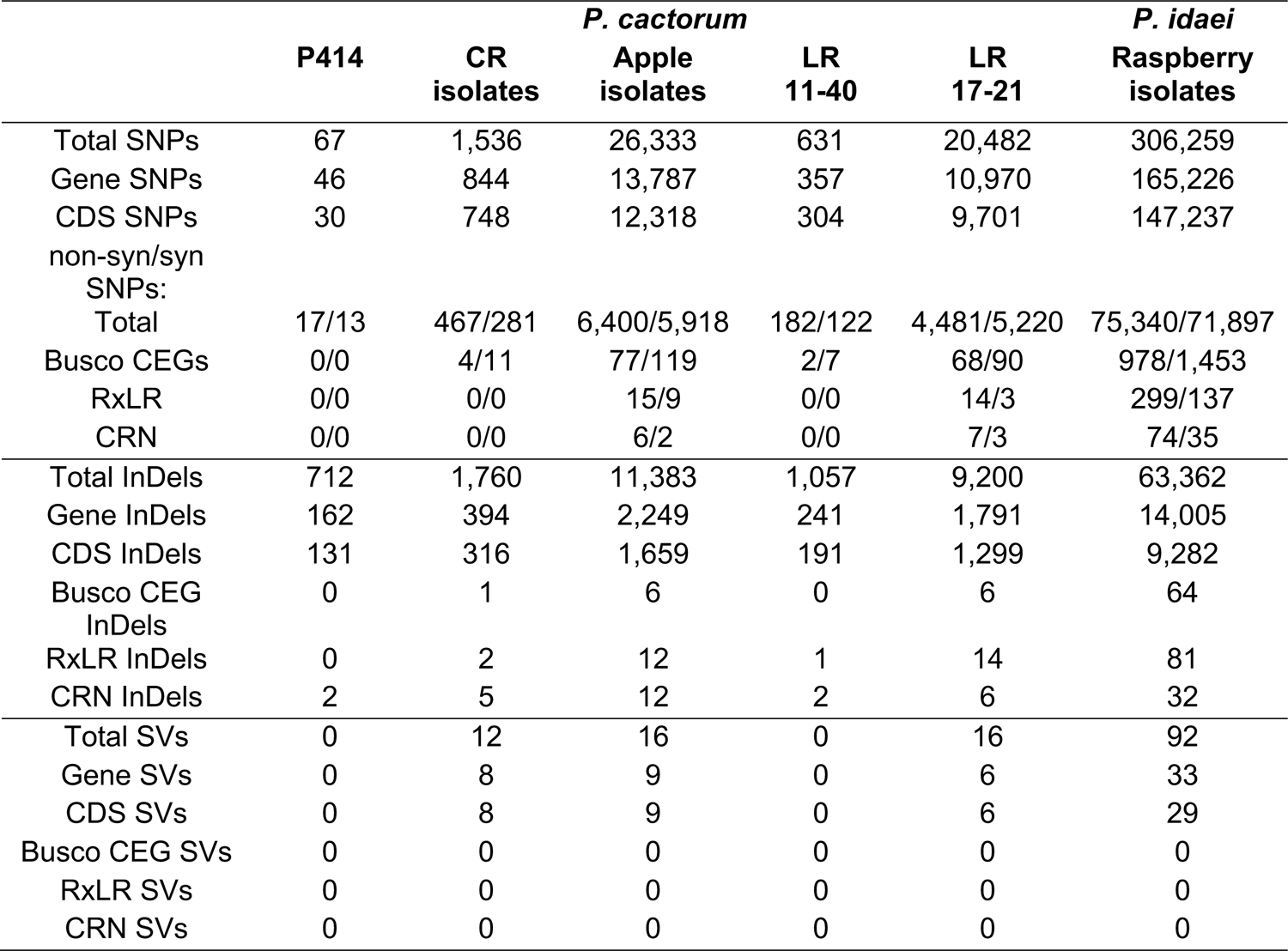
Variant calls vs the reference P414 genome. Numbers of SNPs, insertion/deletion events (InDels) and structural variants predicted in relation to strawberry CR P414. Variants leading to changes within gene models and leading to coding changes in CDS are shown. SNP calls within CDS include the numbers of synonymous/non-synonymous variants.

### DIFFERENTIAL GENE EXPRESSION HIGHLIGHTS KEY EFFECTORS DURING STRAWBERRY INFECTION

#### Putative effectors are highly represented in DEGs during infection of strawberry

RNAseq analysis of *P. cactorum* (P414) infecting strawberry at 12 and 48 hpi showed predicted gene models of putative effectors were upregulated during strawberry infection. Differential gene expression was calculated between mycelium, 12 and 48 hpi in both ‘Emily’ and ‘Fenella’ cultivars and between 12 and 48 hpi timepoints for both cultivars. This allowed identification of early and late expressed transcripts and of the remaining transcripts, identification of those up- and down-regulated *in planta*. In total 9,178 transcripts with LFC >2 were identified. This equated to 34 % of the total transcripts predicted in the genome (**Table 4**). Putative apoplastic and cytoplasmic effectors were overrepresented within the DEGs with 43-76% of candidates differentially expressed in the experiment. Of these, many secreted CAZYme and RxLR candidates showed temporal expression, showing differential expression only the early or late timepoint (160 and 63 transcripts, respectively), with CAZYmes showing a greater number of late expressed candidates and RxLRs showing greater numbers of early expressed candidates (**Table 4** and **Figure 5A**). Furthermore, transglutaminase candidates and Kazal-type protease inhibitors were expressed during early infection, whereas NLP candidates showed a bias towards later infection. Two homologues of *P. infestans* INF1 showed consistent expression across the timepoints (**Supplementary Table 5**; *Pcac1_g22873* and *Pcac1_g22879*). Interestingly, effectors from each category were identified as down-regulated *in planta*, particularly cytoplasmic CRN effectors, of which 69 were down-regulated at both 12 and 48 hpi (**Table 4** and **Figure 5B**). In total, of the 158 putative RxLR effectors identified in P414, just over half, 86 were not expressed or showed low-expression *in planta* (with FPKM values <20) in the RNAseq experiment.

**FIGURE 5.**
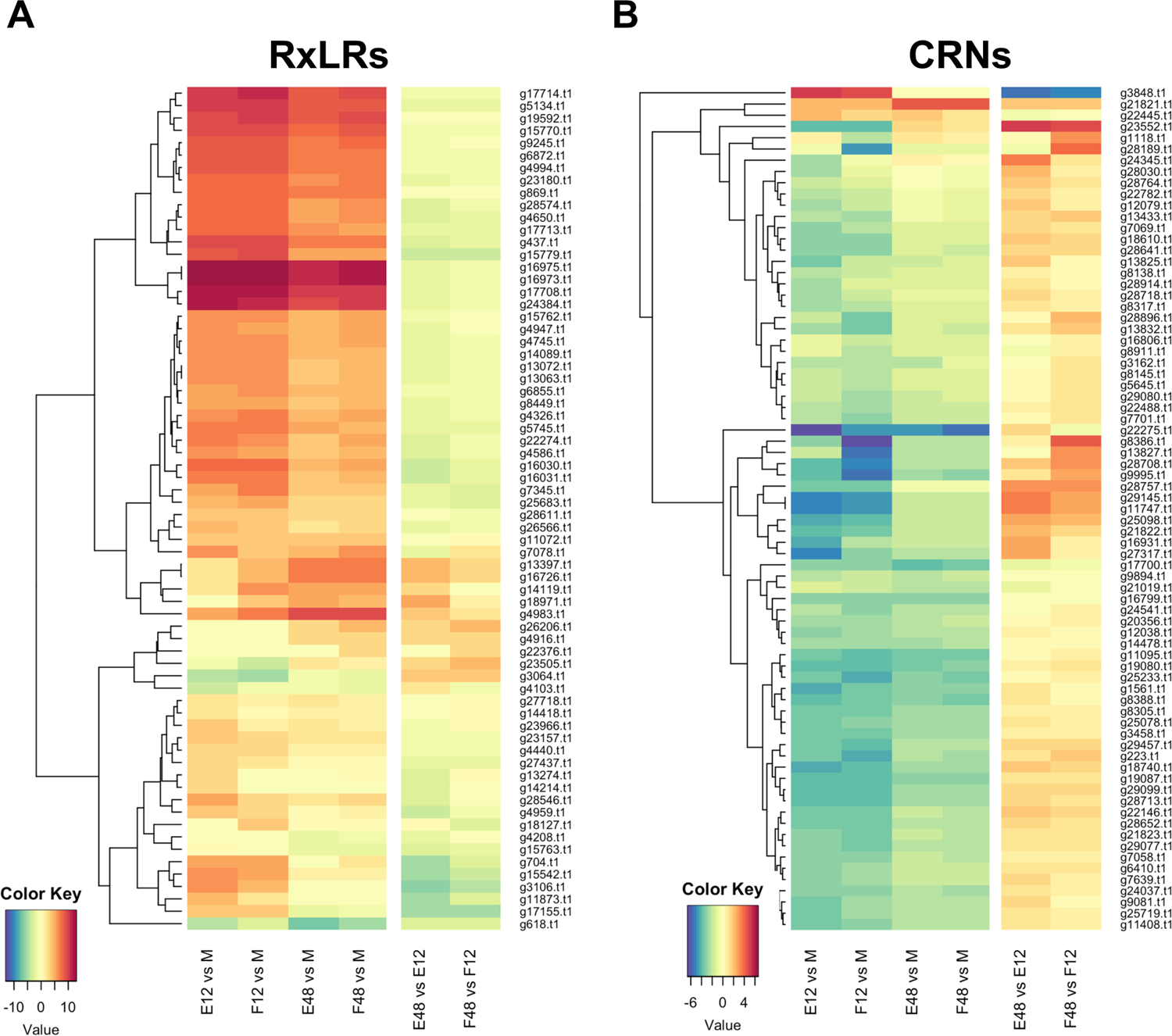
Expression pattern of effectors observed upon infection of both susceptible (‘Emily’) and resistant (‘Fenella’) strawberry roots. RxLR and crinkler (CRN) genes clustered by expression profile. All gene IDs related to the P414 genome (Pcac1_). E: ‘Emily’, F: ‘Fenella’, M: mycelia, 12: 12 hours post inoculation (hpi), 48: 48 hpi. (A) RxLR genes shown in dark red were in the top 25 expressed genes at 12 hpi ranked by log fold change (LFC), (B) the time points investigated don’t appear to have captured differential expression of CRNs.

**TABLE 4.** Expression profile of effector candidates. Numbers of differentially expressed (DE) transcripts, summarised by effector family, showing DE at 24 hours (early induced) or 48 hours (late induced) in both *Fragaria* x *ananassa* ‘Emily’ and ‘Fenella’ cultivars, or upregulated at both timepoints or downregulated at both timepoints *in planta*.

|  | Early induced | Late induced | Upregulated *in planta* | Downregulated *in planta* | Other DE<sup>a</sup> | Total in genome | % DEGs of total<sup>b</sup> |
| --- | --- | --- | --- | --- | --- | --- | --- |
| Total transcripts | 437 | 809 | 2,613 | 5,319 | 1,117 | 29,913 | 34.4 |
| Secreted | 80 | 141 | 442 | 351 | 91 | 1,887 | 58.6 |
| MAMP: Elicitin | 4 | 6 | 23 | 13 | 5 | 67 | 76.1 |
| MAMP: Transglutaminase | 4 | 0 | 4 | 3 | 1 | 18 | 66.7 |
| Apoplastic: Secreted CAZymes | 19 | 36 | 105 | 40 | 14 | 281 | 76.2 |
| Apoplastic: Cutinase | 1 | 0 | 3 | 0 | 0 | 6 | 66.7 |
| Apoplastic: Glucanase inhibitor | 1 | 1 | 10 | 4 | 0 | 30 | 53.3 |
| Apoplastic: NLP | 2 | 6 | 10 | 3 | 0 | 35 | 60.0 |
| Apoplastic: Phytotoxin | 0 | 0 | 1 | 1 | 0 | 3 | 66.7 |
| Apoplastic: Protease inhibitor (cathepsin) | 0 | 0 | 0 | 2 | 1 | 4 | 75.0 |
| Apoplastic: Protease inhibitor (cystatin-like) | 0 | 0 | 2 | 0 | 1 | 5 | 60.0 |
| Apoplastic: Protease inhibitor (Kazal-type) | 5 | 1 | 6 | 4 | 0 | 23 | 69.6 |
| Cytoplasmic: RxLR | 9 | 6 | 48 | 4 | 1 | 158 | 43.0 |
| Cytoplasmic: CRN | 1 | 1 | 2 | 69 | 2 | 127 | 59.1 |

<sup>a</sup> DE - Differential expression.
<sup>b</sup> DEGs – Differentially expressed genes.

#### The most upregulated genes *in planta* include a broad range of effector candidates

Genes involved in initial establishment of infection were further investigated through ranking transcripts by LFC at 12 hpi in comparison to mycelium in the susceptible host ‘Emily’ (all data can be found in **Supplementary Table 5**). In the top 100 ranked genes (LFC values of 16.8-8), secreted proteins were highly represented with 56 present, including 15 RxLR candidates (along with two additional candidates, that carried RxLR motifs but lacked EER motifs), 13 secreted CAZymes (AA7, CBM63, CE8, GH12, GH28 and PL3 families), four NLP candidates, three kazal-type protease inhibitors (one of which was homologous to *P. infestans* EPI10), two glucanase inhibitors and *P. cactorum* factor (PcF) homologue phytotoxin.

Focusing on the top 25 ranked genes (summarised in **Table 5**), six RxLR candidates were upregulated upon infection of strawberry (*Pcac1_g16973*, *Pcac1_g16975*, *Pcac1_g17708*, *Pcac1_g24384*, *Pcac1_g5134* and *Pcac1_g17714*) and an additional one (*Pcac1_g998*), which carried an RxLR motif without an EER motif. *Pcac1_g24384.t1* (hereafter *PcAvh215*) was of particular interest as it was identified as unique to *P. cactorum* strawberry CR isolates and highly expressed *in planta*. Subsequent RT-qPCR analysis in strawberry fruit supported the findings of the RNAseq timepoints and showed expression of *PcAvh215* (**Figure 6A**). Notably, this was the only example of a gene private to strawberry CR isolates in the top 100 ranked transcripts. Investigation into this gene showed it to be a homologue of *P. parasitica* XM_008912329 and *P. sojae Avh32* (JN253712; paralogue to *Avh6;* Wang et al., 2011) with 86% and 82% pairwise amino acid (aa) identity, respectively, downstream of the signal peptide (26-147 aa). *Pcac1_g16973/5*, hereafter *PcAvh136*, was a gene duplication event. *Pcac1_g5134* was the only one out of eight RxLRs with a unique polymorphism (non-synonymous SNP, G53A) between strawberry CR isolates and the three apple and 17-21 isolates, that was expressed.

**FIGURE 6.**
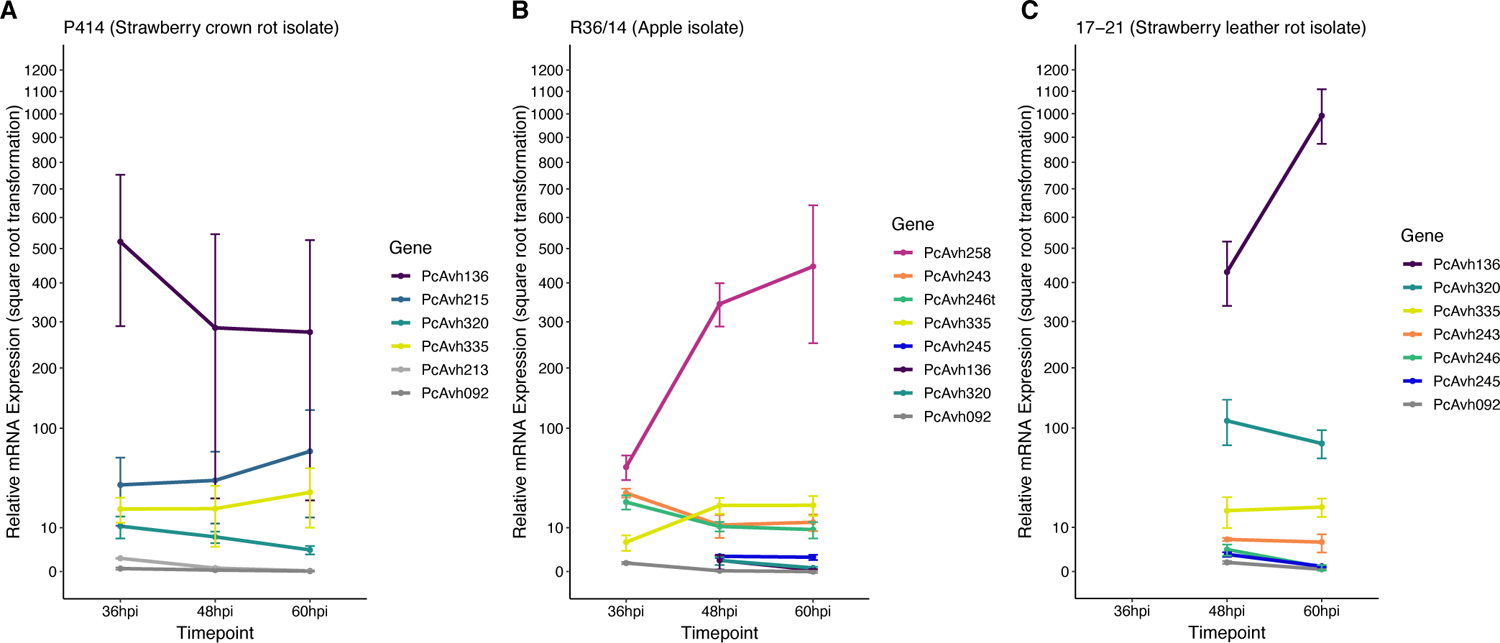
P414, R36/14 and 17-21 RxLR genes show differential expression upon infection of strawberry fruit. (A)-(C) Relative expression of a selection of candidate RxLR genes in three representative Phytophthora cactorum isolates. Gene names listed are for the respective genome. The same gene from different genomes are indicated by the same colour. Genes of interest were normalised to two endogenous reference genes, a ribosomal 40S protein and a protein of the BAR-domain family, Pc_WS41 (Yan and Liou, 2006) and were plotted relative to the expression of the gene of interest in mycelia. Data are the mean of three biological replicates ± se. (A) P. cactorum isolate P414, strawberry crown rot pathotype representative, (B) P. cactorum isolate R36/14, apple pathotype representative, (C) P. cactorum isolate 17-21, strawberry leather rot pathotype representative.

**TABLE 5.** The most upregulated genes in ‘Emily’ at 12 hours post infection include a broad range of effector candidates. Transcripts were ranked by log fold change (LFC) in descending order. Genes private to *Phytophthora cactorum*, with no orthologs in *Phytophthora idaei* were identified, along with those showing single nucleotide polymorphism (SNP) variation at the species, pathotype and population level. Functional annotation of proteins provide evidence for effector candidates.

| LFC | Expression pattern | Transcript ID | Contig | Orthogroup | Presence by group | Presence variation at level: | non-Syn SNP / indel variation at level: | Secretion evidence | Function annotation |
| --- | --- | --- | --- | --- | --- | --- | --- | --- | --- |
| 16.8 | early | g20321.t1 | contig_57 | orthogroup89 | CR(14)LR(1)Md(3)Ri(3) |  |  |  |  |
| 15.6 | early | g2828.t1 | contig_4 | orthogroup15102 | CR(10)LR(1)Md(2)Ri(0) | * |  | Phobius | Coil domain |
| 15.5 | early | g9289.t1 | contig_18 | orthogroup16991 | CR(7)LR(0)Md(0)Ri(0) | * |  |  |  |
| 15.3 | early | g12985.t1 | contig_29 |  |  | * |  |  | Coil domain |
| 14 | early | g7826.t1 | contig_14 | orthogroup15950 | CR(9)LR(0)Md(1)Ri(0) | * | species |  |  |
| 13.6 |  | g12191.t1 | contig_26 | orthogroup359 | CR(14)LR(1)Md(3)Ri(3) |  |  | SignalP; Phobius | EffectorP; Jacalin-like lectin |
| 13.1 |  | g998.t1 | contig_2 | orthogroup774 | CR(14)LR(1)Md(3)Ri(3) |  |  | SignalP; Phobius; TM domain | RxLR (no EER motif); EffectorP |
| 13.1 | early | g24044.t1 | contig_79 | orthogroup14845 | CR(11)LR(1)Md(2)Ri(0) | * | species |  |  |
| 13 |  | g16973.t1 | contig_43 | orthogroup12686 | CR(14)LR(1)Md(3)Ri(0) | *P. cactorum* private |  | SignalP; Phobius | RxLR; EffectorP |
| 13 |  | g16975.t1 | contig_43 | orthogroup12686 | CR(14)LR(1)Md(3)Ri(0) | *P. cactorum* private |  | SignalP; Phobius | RxLR; EffectorP |
| 12.2 |  | g12202.t1 | contig_26 | orthogroup359 | CR(14)LR(1)Md(3)Ri(3) |  | species | SignalP; Phobius | EffectorP; Jacalin-like lectin |
| 11.9 |  | g17708.t1 | contig_46 | orthogroup9680 | CR(14)LR(1)Md(3)Ri(3) |  | species | SignalP; Phobius | RxLR |
| 11.5 |  | g26017.t1 | contig_95 | orthogroup11598 | CR(14)LR(1)Md(3)Ri(3) |  | species | SignalP; Phobius |  |
| 11.4 |  | g24384.t1 | contig_81 | orthogroup14979 | CR(14)LR(0)Md(0)Ri(0) | Strawb. CR private |  | SignalP; Phobius | *PsAvh32* homolog; RxLR; EffectorP |
| 11.2 |  | g20304.t1 | contig_57 | orthogroup947 | CR(14)LR(1)Md(3)Ri(3) |  |  |  | Antibiotic biosynthesis monooxygenase |
| 11.2 |  | g20307.t1 | contig_57 | orthogroup947 | CR(14)LR(1)Md(3)Ri(3) |  |  |  | Antibiotic biosynthesis monooxygenase |
| 11.1 | early | g7647.t1 | contig_14 | orthogroup3422 | CR(14)LR(1)Md(3)Ri(3) |  | species |  | Coil domain |
| 11.1 |  | g9964.t1 | contig_19 | orthogroup13882 | CR(14)LR(1)Md(2)Ri(0) | * | species | SignalP; Phobius | Coil domain |
| 10.8 |  | g613.t1 | contig_1 | orthogroup12809 | CR(14)LR(1)Md(3)Ri(0) | *P. cactorum* private | species | SignalP; Phobius |  |
| 10.7 | early | g4491.t1 | contig_7 | orthogroup201 | CR(14)LR(1)Md(3)Ri(3) |  |  |  | Coil domain |
| 10.7 |  | g13303.t1 | contig_30 | orthogroup8651 | CR(14)LR(1)Md(3)Ri(3) |  | pathotype,species | SignalP; Phobius; TM domain | EffectorP |
| 10.5 |  | g5134.t1 | contig_8 | orthogroup6597 | CR(14)LR(1)Md(3)Ri(3) |  | pathotype,species | SignalP; Phobius; TM domain | RxLR |
| 10.3 |  | g10084.t1 | contig_20 | orthogroup1931 | CR(14)LR(1)Md(3)Ri(3) |  |  | SignalP; Phobius | EffectorP |
| 10.3 | early | g15153.t1 | contig_36 | orthogroup9085 | CR(14)LR(1)Md(3)Ri(3) |  | species | SignalP; Phobius | Protease inhibitor (Kazal-type) |
| 10.3 |  | g15397.t1 | contig_37 | orthogroup1297 | CR(14)LR(1)Md(3)Ri(3) |  | pathotype,population | SignalP; Phobius | Unnamed family (PTHR34737:SF2) |
| 10.3 |  | g17714.t1 | contig_46 | orthogroup9681 | CR(14)LR(1)Md(3)Ri(3) |  |  | SignalP; Phobius | RxLR |

Looking further afield to the remaining nine RxLRs in the top 100 genes, there was another gene duplication event, *Pcac1_g16030/1*, hereafter *PcAvh320*. *PcAvh320* was identified to be a homologue to *PiAvrblb2* (Oh et al., 2009). This homologue was identified as present in all *P. cactorum* isolates. Subsequent RT-qPCR of P414 supported the RNAseq data and showed expression of the gene upon infection of strawberry fruit (**Figure 6A**). *Pcac1_g15770* (hereafter known as *PcAvh266/7/8*) was also identified to be a homologue to another known Avr gene, *PiAvrSmira1* (Rietman et al., 2012).

#### Unique strawberry CR putative secreted proteins are not all expressed during strawberry infection

Of the three putative secreted proteins expanded in strawberry CR isolates, only RxLR candidate *PcAvh215* was found to be differentially expressed by P414 in the time points investigated in strawberry *in vitro* plants (with peak FPKM values of 3,364 and 4,375 in ‘Emily’ and ‘Fenella’, respectively) and strawberry fruit (**Figure 6A**). The other RxLR candidate, *Pcac1_g22827* (hereafter *PcAvh213*), showed low levels of constitutive expression in mycelium with a peak FPKM value of 6 but was not expressed at any time point *in planta* in the RNA-Seq experiment nor subsequent *in planta* RT-qPCR experiment (**Figure 6A**). The uncharacterised secreted protein (*Pcac1_g6287*) had a peak FPKM value of 20 *in planta* but was not investigated in the subsequent RT-qPCR.

### QUANTITATIVE STUDIES OF SELECTED EFFECTORS SHOW DIFFERENTIAL EXPRESSION BETWEEN PATHOTYPE REPRESENTATIVES

#### Candidate RxLR genes show differential expression between isolates from the apple and 17-21 lineage

RT-qPCR analysis of a selection of apple lineage specific (Branch B_0_; **Figure 4**) RxLR candidates show they are expressed by the apple pathotype representative R36/14 upon infection of strawberry fruit (**Figure 6B**), but not the strawberry LR representative isolate 17-21 (**Figure 6C**). The truncated *PcAvh246t* in R36/14 was expressed (**Figure 6B**) but the full-length gene, *PcAvh246*, in 17-21 was not expressed very highly (**Figure 6C**). *PC123_g25079* (hereafter *PcAvh243*) had a similar pattern of expression in R36/14 to *PcAvh246t* but was more highly expressed than *PcAvh246* in 17-21. *PC123_g17462* (hereafter *PcAvh245*) was not expressed very highly in either R36/14 or 17-21 at the time points investigated. *PC123_g15654* was not investigated by RT-qPCR. *PcAvh258*, the only RxLR candidate unique to apple isolates, was the highest expressed RxLR investigated in isolate R36/14 on strawberry fruit (**Figure 6B**).

### Homologues to known avirulence RxLR genes are differentially expressed by the three representative isolates

The expression profiles of a selection of known Avr genes were investigated during strawberry fruit infection by the representative isolates. A homologue to *PiAvramr1* (Lin et al., 2020; *PcAvh335*) was universally expressed in the three isolates investigated (**Figure 6** and **Supplementary Table 6**). In contrast, a homologue to *PiAvramr3* (Lin et al., 2019; *PcAvh092)* was not expressed by any of the isolates at the time points investigated (**Figure 6** and **Supplementary Table 6**). A homologue to *PiAvrblb2* (Oh et al., 2009; *PcAvh320*) was shown to have the greatest upregulation in the strawberry infecting isolates P414 and 17-21, with lower upregulation in R36/14 (**Figure 6** and **Supplementary Table 6**). P414 was predicted to have two copies of the *PiAvrblb2* homologue adjacent to each other on contig 39. Other homologues identified but not investigated with RT-qPCR included *PiAvrvnt1* (Pel, 2010; *PcAvh428/9*) which was found to not be expressed by P414 in the RNAseq data and *PiAvrSmira1* (Rietman et al., 2012; Pc366/7/8) which was found to be expressed *in planta* from the RNAseq data. *PcAvh258*, which was only found in apple isolates, was found to be a homologue to *PiAvr3a*. Other homologues not investigated by RT-qPCR are detailed in **Supplementary Table 6**.

### Candidate RxLRs associated with strawberry infection

Candidate *PcAvh136* was highly expressed by P414 *in planta* and was investigated further in the three representative isolates, where it was found to be the highest expressed RxLR effector investigated in both P414 and 17-21 in strawberry fruit by RT-qPCR. The gene although present in the R36/14 genome was not expressed by the apple isolate, and therefore appears to only be expressed by the two strawberry infecting isolates. Candidate *PcAvh320* appears to be expressed higher by P414 and 17-21 compared to R36/14.

## DISCUSSION

Understanding the pathogenicity of plant pathogens is necessary for designing and implementing durable resistance strategies. Pathogen-host interactions are notoriously dynamic and *Phytophthora* spp. exhibit rapid adaptability to host immunity (Wang and Jiao, 2019; Chepsergon et al., 2020). *P. cactorum* is a continuing threat to strawberry and apple production, as well as more generally to forest trees and woody perennials. Our study revealed that isolates of *P. cactorum* have clear differences in pathogenicity and are generally specialised to different hosts. Sequencing of the contemporary isolates of strawberry CR revealed that there is clear separation between the *P. cactorum* isolates infecting strawberry crowns and apple. We determined that although isolates of *P. cactorum* causing strawberry CR are genetically similar globally, variation in virulence between the isolates was observed, especially on host cultivars with a higher level of resistance. Furthermore, our results highlight several RxLRs that warrant further investigation as host specificity determinants for strawberry and apple.

*P. cactorum* is often described as a generalist pathogen with a wide host range (Grenville-Briggs et al., 2017; Yang et al., 2018). However, our results support previous reports (Seemüller and Schmidle, 1979) which showed that *P. cactorum* isolates originating from strawberry crowns are specialised and less pathogenic on apple bark tissue than isolates originating from strawberry fruit or apple. Conversely Seemüller et al. showed that apple and strawberry fruit isolates were less pathogenic on strawberry crowns. Our data showing the segregation of the strawberry CR and apple isolates into different clades demonstrates clear patterns of separation, which are backed up by analysis of population structure, potentially indicating a species complex and not a single *P. cactorum* species. Of note, the two strawberry LR isolates from our study appear to have a broader host range than either strawberry CR isolates or apple isolates and are able to both cause disease in both strawberry crowns and apple excised shoots but to a more limited extent than most apple isolates. Interestingly, these isolates fall both within the clonal strawberry CR lineages and the more diverse ‘apple’ clade. This may be what has led to confusion in the past, as it is clear that some isolates have a slightly broader host range, but potentially at the expense of enhanced virulence on a single host. However, from our work it is hard to draw conclusions from the strawberry LR isolates as only two isolates were investigated in this study. Further investigation of the isolates sequenced in this study through further RNAseq analysis of infection and the collection of additional isolates may provide a greater understanding of the components required to be a successful pathogen on both strawberry and apple. Screening all isolates on additional potential hosts was not part of the scope of this work but would be an appropriate next step to determine the host range of the isolates.

Clonal pathogen populations are a threat to global food security, as the advantageous stabilising of favourable multi-locus associations results in the rapid spread of specialised lineages within susceptible host populations. These populations can readily adapt to new introduced resistance genes, despite the proportionately moderate levels of standing genetic diversity (Stukenbrock and Bataillon, 2012), presumably due to their high census population sizes (Barton, 2010). In asexual lineages of plant pathogens such as *P. cactorum*, very little is known about the degree to which genetic variation within these lineages influences the virulence profile of populations. We observed a lack of SNP diversity between the strawberry CR isolates, despite isolates being collected from multiple countries, across multiple years, indicating clonality and the possibility of a recent bottleneck. Despite the lack of genetic diversity, substantial variation in pathogenicity on ‘Elsanta’ and ‘Fenella’ strawberry crowns was observed between the strawberry CR isolates. Further investigation is needed to understand whether this variation is either the result of what little genetic variation there is, or whether there is also stably inherited epigenetic silencing of effectors, which has recently been reported in *P. sojae* (Wang et al., 2019) or DNA methylation. N6-methylation (6mA) was profiled in both *P. infestans* and *P. sojae*, however, its exact role in the regulation of gene expression remains to be fully examined (Chen et al., 2018). In the distantly related *P. fragariae*, another pathogen of strawberry, our previous work highlighted that variation in virulence was not related to DNA sequence variation but rather differences in the expression of putative effectors was associated with race structure (Adams et al., 2020). It was proposed that silencing of *Avr* RxLR effectors enables isolates to evade recognition in plants possessing the corresponding *R* gene. The exact mechanism of silencing was not determined during the study, but we postulated that variation was attributed to either control in *trans* or stable forms of epigenetic modifications were regulating gene expression (Adams et al., 2020). Little is known about how filamentous pathogens adapt to host plants by factors other than DNA sequence polymorphism. This study revealed that over half (86) of our effector candidates were not expressed or showed low-expression *in planta*. This indicates that there is pervasive silencing of effectors, highlighting the crucial role of RNAseq data when attempting to determine effector candidates for onward study.

Nonhost resistance (NHR), described roughly as the ability of a plant species to ward off the colonisation of all genotypes of a pathogen species, is poorly understood but often considered the most durable form of resistance, due to the large number of independent resistance mechanisms that are likely acting. Studies in a wide range of pathogens provide evidence for the role of secreted effectors in determining host range (Lee et al., 2017). A classic example is the deletion of a single effector gene, *hopQ1-1* in *Pseudomonas syringae* pv. *tomato*, enabling the pathogen to extend its host range within the Solenaceae to the nonhost plant species, *Nicotiana benthamiana* (Wei et al., 2007). However, non-host resistance ranges from single effectors controlling host range to potentially many tens of effectors. A study of 54 *P. infestans* RxLR effectors in the non-host pepper uncovered that up to 36 could be recognised and led to hypersensitive response in some accessions, suggesting that the recognition of multiple *P. infestans* effectors leads to NHR (Lee et al., 2014). This large number of recognised effectors may be due to the evolutionary differences between wild potato and *Capsicum* species – reported to have split around 20 Ma (Särkinen et al., 2013), which although geographically overlapping have distinct ecological niches (Rumold and Aldenderfer, 2016; Barboza et al., 2019). However, it is possible that *Phytophthora* isolates may carry within their genome many more effectors than they actually express (as we highlight above) and so what matters is which effectors are expressed during infection. What also matters is not just the antigenic potential of the effectors that are expressed, but the function of the effector network in the infection process. It is well known that some effectors are able to mask the avirulence potential of other effectors and therefore evidence for activation of the hypersensitive response alone is also insufficient to assess the functional consequence of a lineage possessing any single effector (Derevnina et al., 2021).

From our work it is not possible to tell how many effectors are causing the observed differences in host range between the clonal CR lineage and isolates infective on apple. The reciprocally incompatible interactions that we observe could be due to effectors either being recognised or the inability of the lineage to efficiently manipulate the host. However, our work did identify candidate RxLR’s which may be good candidates for host specificity determinants on strawberry and apple. Two candidate host-range determinants on strawberry crowns and apple that warrant further investigation are PcAvh215 and PcAvh258, respectively. The strawberry LR isolate, 17-21 does not possess PcAvh215 or PcAvh258 but is pathogenic on both strawberry and apple, to a limited extent compared to isolates isolated from the specific host tissue. This discounts these effectors from being solely responsible for pathogenicity on either host, but they possibly enable the isolates possessing them to have greater virulence on the respective host. Candidate effector PcAvh136 was found to be expressed by both the strawberry CR isolate P414 and the strawberry LR isolate 17-21, but not the apple isolate R36/14 upon infection of strawberry fruit, indicating it may possibly be a determinant for pathogenicity on strawberry. It may also be that PcAvh215 enables isolates possessing it to be more pathogenic on strawberry crowns (all strawberry CR isolates and strawberry LR isolate 11-40), but further RNAseq and transformation of it into non-PcAvh215 containing isolates such as 17-21 would be required to investigate this hypothesis further. Of the effectors that were found to be polymorphic between apple and strawberry only one showed evidence for expression in strawberry.

Understanding effector profiles have aided the characterisation of resistance genes (Armstrong et al., 2005; Poppel et al., 2008; Champouret et al., 2009; Oh et al., 2009; Gilroy et al., 2011; Rietman et al., 2012; Sugimoto et al., 2012), and thereby provided an insight into resistance durability in the field. Homologues to major gene targets (homologues to known *Avr* genes) that are expressed by *P. cactorum* include, *PiAvr1* (Champouret et al., 2009) *PiAvrblb1* (Champouret et al., 2009), *PiAvrblb2* (Oh et al., 2009), *PiAvrSmira1* (Rietman et al., 2012) and *PiAvramr1* (Lin et al., 2020). It would be interesting to investigate if homologues to the resistance genes *Rpiblb1*, *Rpiblb2* and *RpiSmira1* are involved in resistance to *P. cactorum* in strawberry and apple. In addition, there are multiple major gene resistance targets in *P. cactorum* that are present but were not expressed in the *in planta* RNAseq time course of the strawberry CR isolate P414, for example, *PiAvrvnt1* (Pel, 2010), *PiAvr4* (Poppel et al., 2008), *PiAvr8/PiAvrSmira2* (Rietman et al., 2012) and *PiAvramr3* (Lin et al., 2019), it is unknown if these play a role in virulence in *P. cactorum*.

Copy number variation (CNV) is also another form of genetic adaptation that has been identified in *Phytophthora* genomes (Qutob et al., 2009). For example, *P. sojae, P. parasitica* and *P. infestans* genomes encode at least 2, 8 and 11 *Avrblb2* homologues, respectively (Naveed et al., 2019). In the most contiguous *P. cactorum* genome of strawberry CR isolate P414, two homologues, identical to each other (PcAvh320) were predicted adjacent to each other in the genome and were expressed upon infection of strawberry roots. This example of effector duplication therefore supplies the raw materials for adaptive evolution of the gene into novel functions in *P. cactorum*. Further long read sequencing of additional isolates is required to determine the extent of CNV in *P. cactorum* as it is clear that short-read sequencing fails to detect instances of gene duplication with the same fidelity as long-read sequencing.

*In planta* transcriptome sequencing of further isolates will be able to investigate both effector expression variation and the role for CNV’s, alongside functional characterisation and genetic analysis of the basis of resistance to *Phytophthora* in both strawberry and apple. This is non-trivial in both of these crops due to the lack of appropriate methods to transiently induce expression, as infiltration techniques into leaves are challenging due to leaf properties and until extremely recently genome sequences of the octoploid strawberry were not available to aid with the characterisation of putative resistance genes (Edger et al., 2019).

## CONCLUSION

This study provides further evidence that *P. cactorum* should be regarded as a species complex and not a single species, as it comprises of distinct phylogenetic lineages that resolve groups of isolates with distinct effector profiles and displaying host-preference. Pathotype specific effector genes, such as homologues to *PsAvh32* (*PcAvh215*) and *PiAvr3a* (*PcAvh258*) may play roles in specialisation of *P. cactorum* to strawberry and apple, respectively. However, functional analysis is required to validate these genes as determinants of pathogenicity in their respective host.

Further questions that also remain unanswered are to what extent do the expression profiles of effectors in different isolates affect pathogenicity? This highlights the need for further RNAseq from multiple isolates, as well as knockouts to detangle which effectors are differential for pathogenicity in strawberry and apple? This will help us understand what are the key processes that underpin variation in virulence in *P. cactorum*.

This study raises questions about the strategy for effector-informed breeding. We know that clonal lineages predominate in strawberry, as has often been described in other agricultural-associated pathosystems (Hessenauer et al., 2021). We have shown that there are highly expressed lineage-specific effectors within the clonal lineage of CR along with CNV in highly expressed effectors. We hypothesise that these are associated with increased virulence and potentially also pathogenicity itself. If these were then used to screen for and then deploy resistance against them, then the question arises what would the response be, if the pathogen were to adapt? It could be that there are many other effectors that could play a similar role, that are currently silent within the genome, or conversely that through silencing of these effectors, host resistance may be evaded, but at a fitness cost to the pathogen. However, this remains to be seen. Conversely, we see that many ‘core’ effectors that are conserved across lineages on different hosts are present but not expressed, so these, while we may assume they are important from DNA sequence data alone, they are clearly dispensable for pathogenicity, indicating that the idea of core effectors needs to expand beyond simply an analysis of their presence within a genome. Therefore, we must find ways to understand how the network of effectors functions in any given host and whether there are critical effector combinations that if targeted by multiple resistance genes would lead to a durable resistance.

## METHODS

### Phytophthora isolates investigated in this study

Eighteen *P. cactorum* isolates and three *P. idaei* isolates (detailed in **Supplementary Table 1**) were investigated in this study. Of the 18 *P. cactorum* isolates, 13 of these were isolated from strawberry crown tissue exhibiting crown rot symptoms, two from strawberry fruit exhibiting leather rot symptoms and three isolated from symptomatic apple bark. One of the crown rot isolates, 10300, has been previously published by (Armitage et al., 2018). The three isolates of *P. idaei* were isolated from infected raspberry material. All isolates were revived and maintained on V8 agar at 20 °C in the dark.

### Whole genome sequencing and assembly

Mycelia of *P. cactorum* and *P. idaei* were grown in clarified V8-juice broth, similar to (Wilcox et al., 1993); it was comprised of 100 mL V8-juice (Arnotts Biscuits Limited), 1.4 g calcium carbonate (CaCO_3_; Sigma Aldrich) and 100 mL dH_2_O, which were centrifuged at 2,500 x *g* for 15 min, the supernatant was decanted and made up to 1,600 mL with dH_2_O; 200 mL aliquots were dispensed into 250 mL flasks and autoclaved for 20 min at 120 °C. Five mycelial plugs per isolate were added to each flask and were grown at 20 °C under lab light/dark cycle for 10 days in a shaker incubator set to 200 rpm (revolutions per minute). The mycelia were washed in sterile dH_2_O, vacuum filtered and freeze dried overnight.

For one *P. idaei* isolate, SCRP371, gDNA was extracted from freeze dried mycelia using the GenElute Plant Genomic DNA Kit (Sigma). gDNA was sonicated in a water bath and size selected, ∼500 bp, on an agarose gel and extracted. An Illumina library was constructed using the TruSeq LT Kit (FC-121-2001) and was sequenced using Illumina MiSeq v3 2x 300 bp PE Reagent Kit. For all remaining isolates gDNA was extracted from freeze dried mycelia using the Macherey-Nagel Nucleospin Plant II Kit (Fisher Scientific). gDNA was sheared using the Covaris M220 with microTUBE-50 (Covaris) and size selected, 450-600 bp, using a Blue Pippin (Sage Science). Illumina libraries were constructed using a PCR-free method using NEBNext End Repair (E6050S), NEBNext dA-tailing (E6053S) and Blunt T/A ligase (M0367S) New England Biolabs modules. Library insert sizes were 400-600 bp and were sequenced using Illumina Miseq v2 2x 250 bp paired-end (PE; MS-102-2003) or v3 2x 300 bp PE (MS-102-3003) Reagent Kits.

A single *P. cactorum* isolate (P414), isolated from a symptomatic crown of strawberry, was selected for additional PacBio sequencing. gDNA extraction was performed using the Genomic-tip DNA 100/G Kit (Qiagen), following the Tissue Sample method. A minimum of 20 μg of gDNA at approximately 100 ng/μL concentration, with a 260/280 ratio of 1.88 and a 260/230 ratio of 2.26, and a minimum molecular weight of 40 kb was sent to The Earlham Institute, UK. The large insert library was prepared by The Earlham Institute according to manufacturer specifications and sequenced to achieve approximately 87 times coverage on a PacBio RSII platform, using P6-C4 chemistry.

A long-read *de novo* assembly was generated for isolate P414 by first performing read correction and trimming using Canu v1.6 (Koren et al., 2017), before assembling with SMARTdenovo (February 26, 2017 github commit). Errors in this SMARTdenovo assembly were polished through five iterations of Pilon v1.17 (Walker et al., 2014), using the “diploid” flag and trimmed Illumina reads. Illumina reads were trimmed to remove low quality bases and Illumina adapters with fastq-mcf v1.04.676 (Aronesty, 2013).

*De novo* assembly of MiSeq data for the remaining 19 genomes was performed using SPAdes v.3.11.0 (Bankevich et al., 2012). Assembly statistics were collected for all assemblies using QUAST v3.0 (Gurevich et al., 2013). Completeness of the *Phytophthora* genome assemblies was assessed by analysis of conserved Benchmarking Universal Single-Copy Ortholog (BUSCO, v3; (Simão et al., 2015; Waterhouse et al., 2017) genes using the Alveolata-Stramenopiles dataset. DeconSeq was run on all assemblies to remove any potential bacterial contaminants with homology to databases of all ‘complete’ *Bacillus* or *Paenobacillus* genomes as downloaded from NCBI (Schmieder and Edwards, 2011). A database of *Phytophthora* contigs was also made and contigs that showed homology to both bacterial and *Phytophthora* databases were retained. Assemblies were edited in accordance with results from the NCBI contamination screen (run as part of submission to GenBank in December 2017) with contigs split, trimmed or excluded as required. RepeatModeler, RepeatMasker and transposonPSI were used to identify repetitive and low complexity regions (http://www.repeatmasker.org, http://transposonpsi.sourceforge.net).

### Gene and open reading frame prediction

Gene prediction was performed on the softmask*e*d *P. cactorum* genomes using BRAKER1 v.2 (Hoff et al., 2016), a pipeline for automated training and gene prediction of AUGUSTUS v3.1 (Stanke and Morgenstern, 2005). Evidence for gene models were produced using RNAseq data generated as part of this study (discussed below) and aligned to the *P. cactorum* assembly using STAR v2.5.3a (Dobin et al., 2013). Additional gene models were called in intergenic regions using CodingQuarry v.2.0 (Testa et al., 2015), which was run using the “pathogen” flag. Gene models were also augmented with further effector candidates from open reading frames (ORFs) using the methods previously described in (Armitage et al., 2018).

### Functional annotation

Functional annotation of gene models was performed as described previously in (Armitage et al., 2018). Putative CRNs were identified in predicted proteomes and translated ORFs by HMM (Hidden Markov Model) searches using LFLAK and DWL HMM models described in (Armitage et al., 2018). Putative RxLRs were identified following methods previously described by (Armitage et al., 2018). To confirm the absence of candidate genes in our genomes, a Basic Local Alignment Search Tool (BLAST) database was generated in Geneious Prime v2020.0 and used to confirm that genes were not present rather than not being in the respective annotation.

In addition, a publicly available *P. cactorum* genome, LV007, isolated from European Beech (*Fagus sylvatica*) was downloaded from GenBank (PRJNA380728; (Grenville-Briggs et al., 2017) and used in the subsequent analyses (detailed in **Supplementary Table 1**).

### Phylogenetics

A phylogeny was determined from conserved single copy genes present in *P. cactorum* genomes and in *P. idaei* outgroup isolates. Partial and complete single hits from BUSCO searches, using the Alveolata-Stramenopiles obd9 database, were extracted from the 20 sequenced genomes, as well as the publicly available *P. cactorum* 10300 (Armitage et al., 2018) and LV007 genomes (Grenville-Briggs et al., 2017). This led to retention of nucleotide sequences for 230/234 loci, which were aligned using MAFFT v6.864b (Katoh and Standley, 2013), before being trimmed with trimAl v.1.4.1 (Capella-Gutiérrez et al., 2009). A maximum likelihood tree was determined for each locus using RAxML v.8.1.17 (Liu et al., 2011), with the most parsimonious tree for each locus used to determine an overall consensus phylogeny across all 230 loci using ASTRAL v.5.6.1 (Zhang et al., 2018). The resulting tree was visualised using the R package GGtree v.1.12.4 (Yu et al., 2016).

### SNP and variant calling

Single Nucleotide Polymorphisms (SNPs), indels and structural variants were identified in reference to the *P. cactorum* P414 genome. Trimmed Illumina reads from each isolate were aligned to the P414 genome using Bowtie2 v2.2.6 (Langmead and Salzberg, 2012), with SNP variants identified using GATK (McKenna et al., 2010; DePristo et al., 2011; Auwera et al., 2013) and indels/structural variants identified using SvABA (Wala et al., 2018). SNPs called by GATK were filtered using VCFtools (Danecek et al., 2011), retaining bi-allelic SNPs with an QUAL > 30, MQ > 40, DP > 10, GQ > 30. SNP calls were also filtered if isolate P414 Illumina reads showed a homozygous polymorphism in reference to the P414 assembly as these represent errors in the assembly rather than SNP variants. Effects of predicted variants on *P. cactorum* gene models were established using SnpEff (Cingolani et al., 2012). Population genetic statistics were calculated from SNP variants using VCFtools and the R package PopGenome (Pfeifer et al., 2014). Structure analysis was performed using FastSTRUCTURE v1.0 (Raj et al., 2014). The program was run with *k* values between 1-6 and number of populations determined where *k* maximised marginal likelihood. DISTRUCT plots were generated from output meanQ files using R-studio v1.1.453. A SNP distance matrix was made showing the number of variants that differ between isolates. SNP variants were extracted from the final .vcf file as a fasta alignment of concatenated variable sites, containing two sequences per isolate (representing the first and second allele called at each site, respectively). A distance matrix was calculated in Geneious Prime v2020.0 and exported into Microsoft Excel.

### Phytophthora *spp.* zoospore production

The production of zoospores was followed from (Nellist et al., 2019). To summarise, 10 mm discs were excised from the edge of actively growing colonies, the plugs were covered with dilute compost extract in 90 mm triple-vented petri dishes (five per plate; Thermo Scientific) and were placed under constant light conditions at 20 °C, for 48 hours, to stimulate sporangia development. After 48 hours, the diluted compost extract was poured off and replaced with a fresh solution. The plates were moved to a fridge (∼4 °C) and chilled for 45 min and then moved to the bench and left to warm up at room temperature for 45 min, to stimulate the release of zoospores. The solution was then vacuum filtered through Whatman 113V Wet Strengthened 150 mm filter paper and the concentration of zoospores was determined using a haemocytometer and adjusted to 1 x 10^4^, 2 x 10^4^ or 5 x 10^3^ zoospores per mL by diluting with dilute compost extract. The adjusted solution was kept on ice until ready to be used to inoculate plants/unripe fruit.

### Pathogenicity tests on strawberry crowns

The virulence of the 18 *P. cactorum* and three *P. ideai* isolates were tested on the crowns of three cultivars of cultivated strawberry (*F*. x *ananassa*). ‘Malling Opal’ is an everbearer that is extremely susceptible to *P. cactorum*, bred by NIAB EMR (formally East Malling Research) and released in 2005. ‘Elsanta’ is a mid-season variety with reported susceptibility to crown rot. ‘Fenella’ is a mid-late season variety with good resistance to *P. cactorum*, bred by NIAB EMR and released in 2009.

Mother stock plants of the three cultivars were maintained in 1 L pots (Soparco) in polytunnels. Ten runners of each of the three cultivars were pinned down for each isolate into 9 cm diameter pots (Soparco) filled with peat-based compost. The clones were grown on for four and a half months and then placed in a 2 °C coldstore for one week and then moved into a −2 °C coldstore for at least two months. After two months the plants were removed from the coldstore and the dead leaves were removed. The plants were grown on for three weeks in a glasshouse compartment maintained at 20 °C during the day and 15 °C at night, on a 16/8 hour, day/night light cycle, as described by Nellist et al. (2019).

The inoculation procedure for coldstored strawberry plants was performed as described in Nellist et al. (2019). Pathogenicity screens were performed under controlled conditions in glasshouse compartments, maintained at 20 °C during the day and 15 °C at night on a 16/8 day/night cycle for four weeks. Fifteen mm wounds were made at the base of a single petiole using a scalpel and ∼5 mL of 1 x 10^4^ zoospore suspension was sprayed across each wound and onto the compost. Plants were covered with clear plastic sheeting for 48 hours to maintain humidity. Scoring of symptoms was as described by Nellist et al. (2019) based upon a modified version from (Bell et al., 1997), where plants were scored on a scale of 1-8. Foliage was assessed visually for the presence of wilting symptoms, weekly over the four week period. If the plant died during the first, second, third or fourth week after inoculation, it was assigned a score of 8, 7, 6 or 5, respectively. The strawberry crowns were then cut open longitudinally and assessed on a scale of 1-5; 1 - healthy (0% infection), 2 - up to 25% infection, 3 - 26-50% infection, 4 - 51-75% infection, 5 - 76-100% infection. The data for the ten replicates were averaged and a mean crown rot disease score was used for further analysis. Statistical analyses were performed using R (v3.6.0, “Planting of a Tree” (Team, 2019).

### Pathogenicity tests on detached unripe strawberry fruit

Unripe ‘Elsanta’ strawberry fruit were picked while still white/green in the Summer of 2018. The fruit were then surface sterilised by immersion in a 10% bleach solution and then rinsed twice with dH_2_O. The fruit were dried off and two fruit were placed into each sterile 90 mm triple-vented petri dish bottom or lid (Thermo Scientific). The petri dish lids and bottoms with fruit were placed on trays sterilised with 70% ethanol. A sterile 4 mm cork borer was used to bore a shallow hole in the fruit. Zoospores were produced as described above and 100 μL of 5 x 10^3^ zoospore suspension was added into the hole of the fruit. Fruit were screened in three separate experiments with a minimum of eight replicates per isolate screened in experiment. The trays were then sealed in a plastic bag and left in the dark at a constant temperature of 20 °C. The ratios of colonised to non-colonised fruit were recorded after seven days.

### Pathogenicity tests on excised apple shoots

Dormant first year growth apple shoots were collected from ‘Cox’ and ‘Gala’ in the Winter 2018. The shoots were cut to a length of 22 cm and surface sterilised for 15 minutes, by immersing them in a 10% bleach solution. The shoots were then rinsed three times with sterile distilled water and one centimeter was excised from each end. Both ends were sealed by dipping in molten paraffin wax. A wound was produced in the middle of each shoot using a 4 mm diameter cork borer and the outer bark was removed with a scalpel. Agar plugs of the same diameter, containing the leading edge of *P. cactorum* mycelium were placed mycelium-side down onto the wound to inoculate the shoots. Six shoots of each cultivar were inoculated per isolate. Mock inoculation of six shoots per cultivar were performed using sterile V8 agar plugs. The excised shoots were then transferred to boxes, damp paper towels were placed at the bottom of each box and the shoots were randomised by isolate and placed on raised racks made of aluminium foil. The boxes were sealed in clear plastic bags to maintain humidity and were placed in a controlled environment room, with a constant temperature of 22 °C (±2 °C) and a 16/8 hour, light/dark cycle for four weeks. Shoots were assessed for maximum lesion length at four weeks by removing the bark around the wound using a scalpel. A digital caliper was used to take measurements and the original wound size, 4 mm, was subtracted from each measurement.

### In vitro strawberry root pathogenicity transcriptome analysis

Transcriptome changes during host infection were investigated through an infection time-course on strawberry roots infected with *P. cactorum* isolate P414. The time-course was performed in the susceptible cultivar ‘Emily’ and moderately resistant cultivar ‘Fenella’; parents of a mapping population used in a previous study (Nellist et al., 2019). Micropropagated plants were produced by GenTech Propagation Ltd. for these experiments. Upon arrival at NIAB EMR, plants were transferred to 120 x 120 x 15 mm, four vent, petri dishes (Corning, Gosselin), half filled with ATS (*Arabidopsis thaliana* salts) media, two plants per plate. ATS media was prepared as described by (Taylor et al., 2016). The media was poured into the bottom plate and after it had set, half of the agar was excised with a sterile flat spatula. Plants were then transplanted so the crown sat on the top of the agar (**Supplementary Figure 1A**). The roots were gently smoothed down, ensuring they were touching the agar. The plates were then sealed with Sellotape and aluminium foil cases were made to surround the agar ensuring a dark environment for the root system (**Supplementary Figure 1B**). The plates were then positioned upright in a growth cabinet (Panasonic MLR-325H) at 22 °C, on a 16/8 hour, day/night light cycle with a photosynthetic photon flux (PPF) of 150 µmol m^!“^s^!#^provided by fluorescent lamps (FL40SSENW37).

Just before inoculation, the micropropagted ‘Emily’ and ‘Fenella’ plants were transferred to fresh plates and zoospores were produced as described above. Each root system was inoculated with 1 mL of 2 x 10^4^ zoospore suspension, using a pipette and slowly dripping the suspension over the entire root system. The plates were then sealed with Sellotape, partially recovered with the aluminium foil and were kept flat for 2 hours to allow the zoospores to encyst. Mock inoculated (0 hour post inoculation) were inoculated with 1 mL of dilute compost extract. The plates were then returned to their upright position until harvested. Root samples were collected at 0 (mock), 6, 12, 24, 48, 72, 96, 120 and 144 hours post inoculation (hpi). The root systems were swilled in sterile dH_2_O to remove any agar, patted dry and were collected in 2 mL Eppendorf microcentrifuge tubes, flash frozen in liquid nitrogen and stored at −80 °C.

Total RNA was extracted from the strawberry roots following a modified version of Yu et al. (2012), over two days. All solutions were made using RNase-free reagents and water. Mortar and pestles were decontaminated with RNAseZap^TM^ (Invitrogen/Thermo FIsher Scientific) and were baked for two hours at 230 °C. Frozen root material and PVPP (0.01 g:0.1 g of frozen material) were weighed and ground with a mortar and pestle, in the presence of liquid nitrogen. The ground plant material was then split in half and transferred to 2x 700 μL of prewarmed (65 °C) extraction buffer (3% CTAB, 100 mM Tris-HCL pH 8.0, 1.4 M NaCl, 20 mM EDTA, 5% PVP and RNase-free H_2_O) with 10 μL of β-mercaptoethanol. The remaining steps were then performed as previously described by (Yu et al., 2012) and the RNA was eluted into 65 μL DEPC-treated H_2_O. The quantity and purity of the RNA were determined using the NanoDrop (ND-1000, Thermo Fisher Scientific) spectrophotometer. The Qubit 2 Fluorometer (Thermo Fisher Scientific) was also used to more accurately determine RNA quantity. RNA Integrity Number (RIN) was assessed by running the RNA on the Tapestation 4200 (Agilent Technologies). At least 1 μg of root RNA with a RIN score above 7 and with 260/280 and 260/230 ratios above 1.8 were sent to Novogene for sequencing. Strawberry root samples were sequenced to a depth of 50 million reads per sample.

Timepoints for sequencing were selected through the detection of β-tubulin transcripts by Reverse Transcriptase-PCR, using SuperScript^TM^ III Reverse Transcriptase kit with an equal amount of RNA used for each sample. The complementary DNA (cDNA) was then analysed by PCR with 200 μM dNTPs, 0.2 μM of each primer (detailed in **Supplementary Table 2**), 2 μL of cDNA template and 2.5 units of Taq DNA polymerase and the buffer supplied in a 20 μL reaction. Reactions were conducted in a Veriti 96-well thermocycler with an initial denaturation step at 95 °C for 30 s, followed by 35 cycles of a denaturation step at 95 °C for 30 s, an annealing temperature of 60 °C for 30 s and an extension step of 72 °C for 30s. This was followed by a final extension step of 72 °C for 5 min and held at 10 °C. Products were visualised by gel electrophoresis on a 1% w/v agarose gel at 80 V for 90 min, stained with GelRed. Following this, three biological replicates of samples taken at: 0, 12 and 48 hpi for both ‘Emily’ and ‘Fenella’ were sequenced (**Supplementary Table 3**).

Mycelia of P414 were grown in clarified V8-juice broth as described as above with the addition of 500 μg/mL of ampicillin (Fisher) and 10 μg/mL of rifampicin (Fisher) at 20 °C under lab light/dark cycle for 10 days in a shaker incubator set to 200 rpm. The mycelia were washed in sterile dH_2_O, vacuum filtered, flash frozen in liquid nitrogen and stored at −80 °C. Total RNA was extracted from three biological replicates of flash frozen P414 mycelia using the RNeasy Mini Kit (Qiagen) following the manufacturer’s protocol. RNA quality and quantity were assessed as described above and the RNA was sent to The Earlham Institute, UK for sequencing. cDNA library insert sizes were 450-625 bp and were sequenced on Illumina HiSeq4000, using 2x 150 bp PE Reagent Kit. Three barcoded biological replicates of each treatment were pooled and sequenced across multiple lanes. *P. cactorum* mycelia were sequenced to a depth of 25 million reads per sample.

Illumina adapters and low-quality bases were trimmed using fastq-mcf. All RNAseq data was aligned to the *Fragaria vesca* genome v1.1 (Shulaev et al., 2011) using STAR v2.5.3a (Dobin et al., 2013), to remove strawberry reads from the dataset. Read alignment for differential gene expression was performed using Salmon v0.9.1 (Patro et al., 2017) with differential gene expression during infection investigated using DESeq2 (Love et al., 2014). The normalised expression value was represented by applying Fragments per Kilobase of exon model per Million mapped reads (FPKM). All mycelial genes with an FPKM value <1 were adjusted to 1. The Log Fold Change (LFC) in gene expression was calculated using adjusted FPKM values to prevent overprediction of LFC from low-expressed/unexpressed genes under one condition, with all genes with an FPKM value <1 were adjusted 1; LFC = log2(‘Emily’ 12 hpi FPKM)/log2(Adjusted Mycelium FPKM). Genes were designated as differentially expressed if they had a DeSeq2 P-adj <0.05 and LFC was ≯2 or was ≮-2., The top 100 expressed genes were investigated further, based on LFC in descending order of ‘Emily’ at 12 hpi vs Mycelium. Temporal expression was assessed by identifying those differentially expressed genes with a consistent in peak expression (based upon LFC) in both Emily and Fenella at 12hpi (early-expressed genes) or 48hpi (late-expressed genes).

### Comparative genomics of known virulence related factors

A selection of known RxLR *Avr* gene sequences (amino acid sequence after the signal peptide) were BLASTed against the 22 *Phytophthora* genomes in Geneious Prime v2020.0 and the hits (tBLASTx E-value > 1×10^-10^) were investigated. Expression of interesting hits were analysed using the *in planta* RNAseq data and interesting candidates were further explored in the representative isolates in the unripe fruit assay.

### Strawberry fruit Reverse Transcription quantitative Polymerase Chain Reaction (RT-qPCR) screen

Unripe, green/white ‘Driscoll® Amesti™’ strawberry fruit were used for the pathogenicity time course of three *P. cactorum* isolates; P414, R36/14 and 17-21. The fruit were sterilised and prepared as described above. Zoospores were produced as described above and 100 μL of 5 x 10^3^ zoospore suspension was added into the hole of each fruit. Samples were taken at 0, 36, 48 and 60 hpi. A larger cork borer of 10 mm was used to excise an area around the inoculation point of the fruit, the excised samples were flash frozen in liquid nitrogen and stored at −80 °C.

Mycelia of P414, R36/14 and 17-21 were grown in clarified V8-juice broth and harvested as described above. Total RNA was extracted from the fruit as described above for strawberry roots and from the flash frozen mycelia of P414, R36/14 and 17-21 as described above. The RNA was assessed by the NanoDrop and Qubit 2 as described above. RNA samples were normalised to 720 ng. Reverse transcription was performed on three biological replicates of each *in planta* time point and two biological replicates for the mycelia timepoints with the QuantiTech Reverse Transcription Kit (Qiagen). *In planta* timepoints, for further analysis, were confirmed through the positive identification of *P. cactorum* β-tubulin product by Reverse Transcription-Polymerase Chain Reaction (RT-PCR).

Reverse Transcription quantitative PCR (RT-qPCR) was then performed in a CFX96^TM^ Real-Time PCR detection system (BioRad) in 10 μL reactions of: 5 μL of 2x qPCRBIO SyGreen Mix Lo-Rox (PCR Biosystems), 2 μL of a 1:5 dilution of the cDNA sample in dH_2_O and 400 nM of each primer (detailed in **Supplementary Table 2**). Due to the presence of primer dimers, an additional step at the end of the reaction was added to measure the fluorescence at a temperature greater than the melting temperature of the primer dimers (Ball et al., 2003). The reaction was run with the following conditions: 95°C for 2 minutes, 40 cycles of 95°C for 5 seconds, 62°C for 20 seconds and 80°C for 5 seconds. This was followed by 95°C for 10 seconds, and a 5 second step ranging from 65°C to 95°C by 0.5°C every cycle to generate melt curves. At least two technical replicates for each sample were performed and the melt curves were analysed to ensure the correct product was detected. Relative gene expression was calculated using the efficiency corrected method, which determines the relative gene expression ratio based on the real-time PCR efficiencies and the cycle quantification value (C_q_; Pfaffl, 2001):

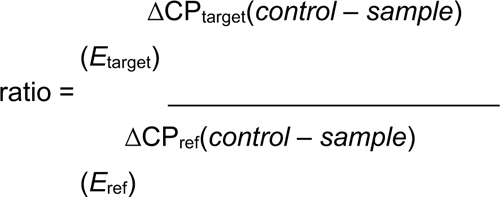

Expression values were calculated as the mean of the three biological replicates and the standard error of the mean was calculated and plotted. A pooled sample of all cDNA was used as an inter-plate control (IPC) on all plates using primers for β-tubulin (Pcac1_g23639; **Supplementary Table 2**). Genes of interest were normalised to two endogenous reference genes (**Supplementary Table 2**), a ribosomal 40S protein (Pcac1_g24902) and a protein of the BAR-domain family, Pc_WS41, Pcac1_g27577 (Yan and Liou, 2006) and were plotted relative to the expression of the gene of interest in mycelia.

## AUTHOR CONTRIBUTIONS

RJH, CFN and ADA devised the study. CFN performed the experimental work, with input from LAL, HJB and ML. ADA performed the bioinformatic analyses including genome assembly, annotation, orthology gene expression and variant calling analyses with input from MKS. RJH, CFN and ADA conceived and drafted the manuscript. All authors read and approved the submission.

## ACKNOWLEDGEMENTS

This research was supported by grants awarded to RJH from the Biotechnology and Biological Sciences Research Council (BBSRC - BB/K017071/1, BB/K017071/2 and BB/N006682/1) and ML who is receipt of PhD funding from the BBSRC Collaborative Training Partnership for Crop Research (CTP_FCR2018-5, AHDB Grant ST/TF170). The authors gratefully acknowledge the NIAB EMR Plant Clinic, Prof. May Bente Brurberg, Dr Thijs van Dijk, Dr Vance M. Whitaker and Prof. Natalia A. Peres for providing *Phytophthora cactorum* isolates. The authors also thank Dr Thomas M. Adams for useful discussions and the East Malling Strawberry Breeding Club for access to strawberry material. In addition, we wish to thank Mr Adam B. Whitehouse, Mr Andrew J. Passey, Mr César Marina-Montes and Mr Joseph Hutchings for the help received in the preparation and setting up of experiments. Work was carried out under the terms of DEFRA Plant Health Licence 6996/221427 held by RJH/CFN.

## SUPPLEMENTARY DATA

SUPPLEMENTARY TABLE 1 | Summary of *Phytophthora cactorum* and *Phytophthora idaei* isolates used in this study.

SUPPLEMENTARY TABLE 2 | Characteristics of primers used in this study.

SUPPLEMENTARY TABLE 3 | Details of genes expanded and contracted in the *Phytophthora cactorum* strawberry crown rot and apple lineages.

SUPPLEMENTARY TABLE 4 | Summary of identified *Phytophthora cactorum* Avirulence gene homologs.

SUPPLEMENTARY TABLE 5 | *Phytophthora cactorum* isolate P414 annotation table and *in planta* RNAseq analysis from *Fragaria* x *ananassa*.

SUPPLEMENTARY TABLE 6 | Summary of homologues to known avirulence RxLR genes in Phytophthora cactorum.

---

**SUPPLEMENTARY FIGURE 1.**
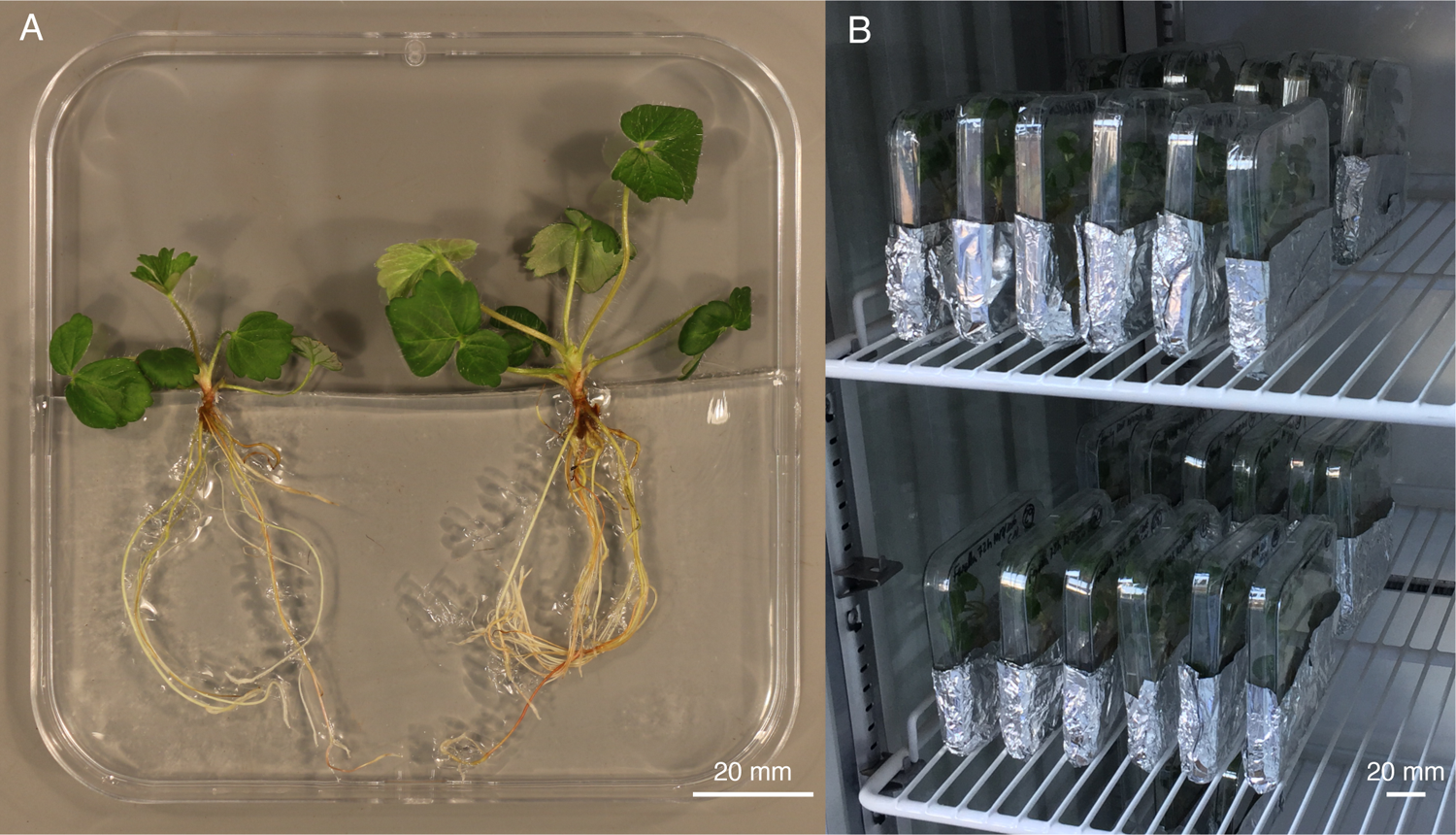
Example experimental setup of *in vitro Fragaria* x *ananassa* plants. **(A)** strawberry plants positioned in petri dishes, on top of agar, **(B)** aluminium foil casing and upright positioning of plates in the growth incubator.

**SUPPLEMENTARY FIGURE 2.**
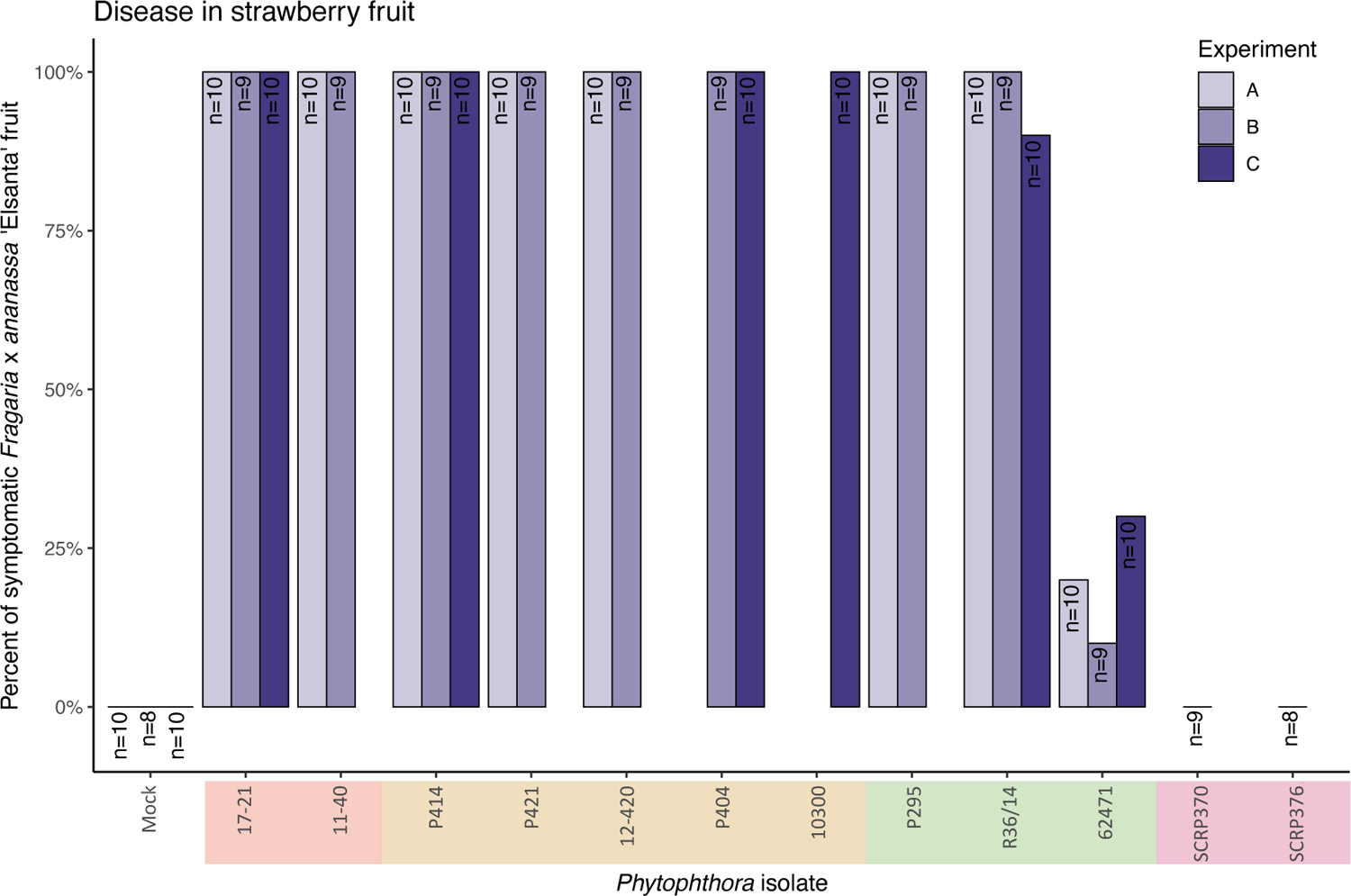
All *Phytophthora cactorum* isolates tested were able to cause disease in strawberry fruit. Percentage of symptomatic strawberry ‘Elsanta’ fruit after artificial inoculation *Phytophthora cactorum* and *Phytophthora idaei* zoospores, from three separate experiments.

**SUPPLEMENTARY FIGURE 3.**
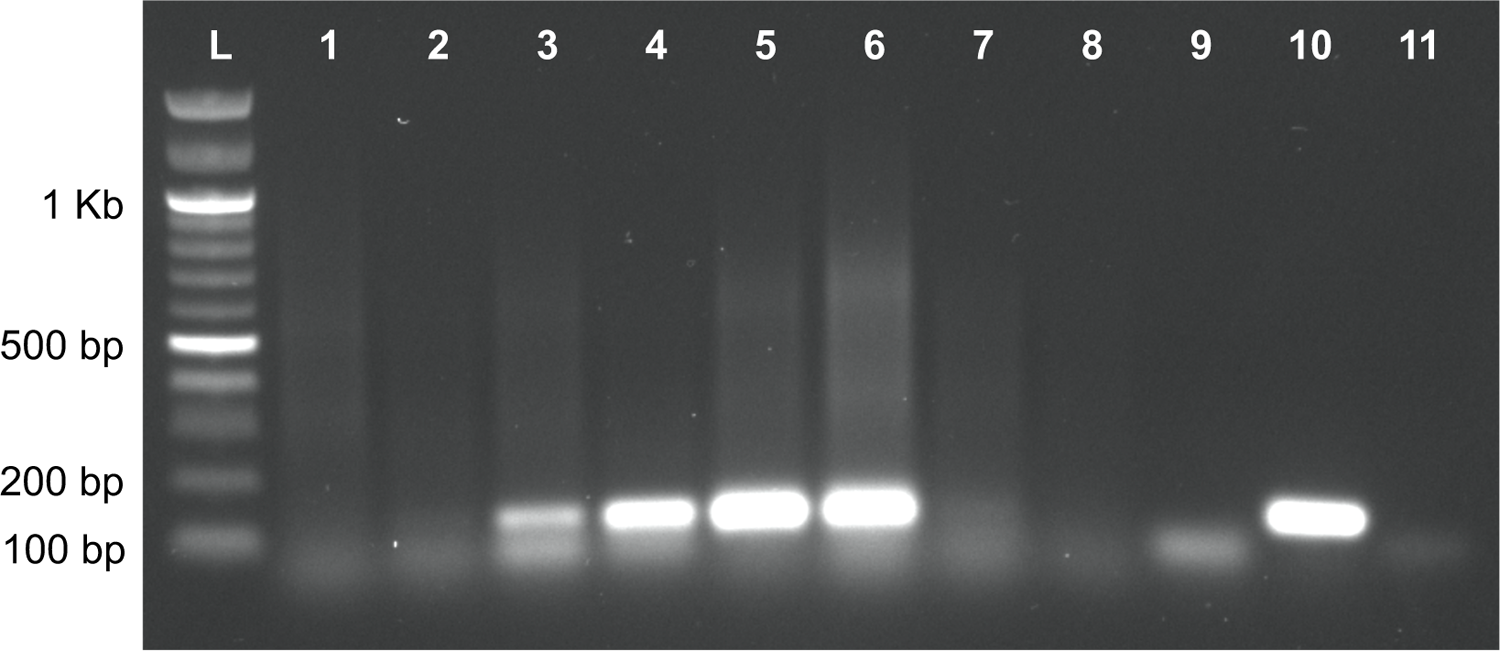
**Agarose gel electrophoresis of RT-PCR β-tubulin reactions on representative samples from inoculation time course experiment of *Phytophthora cactorum* isolate P414 on the ‘Emily’ cultivar of *Fragaria* x *ananassa*. L**: 100 bp DNA Ladder (New England Biolabs). **1**: Mock inoculated ‘Emily’. **2**: 6 hours post inoculation (hpi). **3**: 12 hpi. **4**: 24 hpi. **5**: 48 hpi. **6**: 72 hpi. **7**: 96 hpi. **8**: 120 hpi. **9**: gDNA from ‘Emily’ (negative control). **10**: gDNA from P414 mycelium (positive control). **11**: dH_2_O (negative control).

