## Supplementary material for "Comparative analysis of host-associated variation in *Phytophthora cactorum*": Supp. Table 1 Isolate details.docx

| **Isolate ID** | **Genome ID** | **GenBank Accession** | **Species** | **Material isolated from** | **Year** | **Location** | **Previously published** |
| --- | --- | --- | --- | --- | --- | --- | --- |
| P404 | PC116 | RCMJ00000000 | *P. cactorum* | Strawberry crown | 1998 | UK |  |
| P414 | Pcac1 | NHQK00000000 | *P. cactorum* | Strawberry crown | 2011 | Somerset, UK |  |
| P415 | PC118 | RCML00000000 | *P. cactorum* | Strawberry crown | 2013 | UK |  |
| P416 | PC119 | RCMM00000000 | *P. cactorum* | Strawberry crown |  | UK |  |
| P421 | PC129 | RCMV00000000 | *P. cactorum* | Strawberry crown | 2017 | UK |  |
| PC13/15 | PC122 | RCMP00000000 | *P. cactorum* | Strawberry crown | 2015 | UK |  |
| 10300 | PC110 | GCA_003287315.1 | *P. cactorum* | Strawberry crown | 2006 | Norway | Armitage et al. 2018 |
| 4032 | PC115 | RCMI00000000 | *P. cactorum* | Strawberry crown |  | Netherlands |  |
| 4040 | PC117 | RCMK00000000 | *P. cactorum* | Strawberry crown |  | Netherlands |  |
| 2003-3 | PC114 | RCMH00000000 | *P. cactorum* | Strawberry crown |  | Netherlands |  |
| 12-420 | PC111 | RCME00000000 | *P. cactorum* | Strawberry crown | 2012 | Florida, USA |  |
| 15-7 | PC113 | RCMG00000000 | *P. cactorum* | Strawberry crown | 2015 | Florida, USA |  |
| 15-13 | PC112 | RCMF00000000 | *P. cactorum* | Strawberry crown | 2015 | Florida, USA |  |
| 11-40 | PC127 | RCMT00000000 | *P. cactorum* | Strawberry fruit | 2011 | Florida, USA |  |
| 17-21 | PC128 | RCMU00000000 | *P. cactorum* | Strawberry fruit | 2017 | Florida, USA |  |
| R36/14 | PC123 | RCMQ00000000 | *P. cactorum* | Apple | 2014 | Kent, UK |  |
| 62471 | PC120 | RCMN00000000 | *P. cactorum* | Apple | 2014 | Kent, UK |  |
| P295 | PC121 | RCMO00000000 | *P. cactorum* | Apple (collar rot) | 1984 | Offham, UK |  |
| LV007^a^ |  | GCA_002081965.1 | *P. cactorum* | European Beech | 2016 | Sweden | Grenville-Briggs et al. 2017 |
| SCRP370 | PI125 | RCMR00000000 | *P. idaei* | Raspberry | 1985 | Scotland, UK |  |
| SCRP371 | PI124 | QOKR00000000 | *P. idaei* | Raspberry | 1986 | England, UK |  |
| SCRP376 | PI126 | RCMS00000000 | *P. idaei* | Raspberry | 1993 | England, UK |  |

^a^ Only sequence data was used in this study.
