## Supplementary material for "Comparative analysis of host-associated variation in *Phytophthora cactorum*": Supp. Table 2 Primer Details.docx

| **Primer ID** | ***Avr* homolog** | ***PcAvh ID*** | **P414  gene ID**  **(Pcac1_)** | **R36/14 gene ID**  **(PC123_)** | **17-21 gene ID**  **(PC128_)** | **Sequence (5’-3’)** | **Length** | **Orientation** | **Product size** | **Target region** |
| --- | --- | --- | --- | --- | --- | --- | --- | --- | --- | --- |
| B_tub_F | -- | -- | g23639 | contig 36 | g2365 | AGGAGATGTTCAAGCGTGTG | 20 | Forward | 129 | ß-tubulin |
| B_tub_R2 |  |  |  |  |  | GATCGTTCATGTTGGACTCA | 20 | Reverse |  |  |
| B_tub_F3 | -- | -- | g23639 | contig 36 | g2365 | GATCCCCAACAACATCAAGGCTAG | 24 | Forward | 193 | ß-tubulin |
| B_tub_R3 |  |  |  |  |  | GAACTCCATCTCGTCCATACCTTC | 24 | Reverse |  |  |
| 40S_F1 | -- | -- | g24902 | g6647 | g9173 | GATCGACATCTACATCCCGCG | 21 | Forward | 157 | Ribosomal 40S |
| 40S_R1 |  |  |  |  |  | GTTGATCTGCACCGAGGC | 18 | Reverse |  |  |
| Pc_WS41_F2 | -- | -- | g27577 | g12879 | g9492 | CGGTCCAATCGTGCTCTATTTG | 22 | Forward | 82 | BAR-domain family |
| Pc_WS41_R2 |  |  |  |  |  | ACTGGACTTGAAGGAGAAGCG | 21 | Reverse |  |  |
| PcRx006_F2 | *Pi Avr3a* HQ | *PcAvh258* | -- | g19522 | -- | CAGACTTCGACCAAACCAAGG | 21 | Forward | 207 | RxLR |
| PcRx006_R2 |  |  |  |  |  | CTTCCGCCACCTTATGATTGTC | 22 | Reverse |  |  |
| PcRx008_F2 | -- | *PcAvh213* | g22827 | contig 271 | contig 345 | TACAGTCGCCACCCTCATC | 19 | Forward | 165 | RxLR |
| PcRx008_R3 |  |  |  |  |  | CTTCCGATGCCTCGCTCTTC | 20 | Reverse |  |  |
| PcRx009_F5 | *Ps Avh32* | *PcAvh215* | g24384 | -- | -- | GGGTGACGAAGGCAAAATTACC | 22 | Forward | 191 | RxLR |
| PcRx009_R6 |  |  |  |  |  | CAATGAGACTATCTGGCACACG | 22 | Reverse |  |  |
| PcRx010_F2 | -- | *PcAvh245* | -- | g17462 | g20971 | CGTTGACCTCGCACCAAAT | 19 | Forward | 164 | RxLR |
| PcRx010_R2 |  |  |  |  |  | AGGCATTCCCCTCTCTTCAGT | 21 | Reverse |  |  |
| PcRx011_F2 | -- | *PcAvh243* | -- | g25079 | g19804 | CTTCTGCGGGCTAGTTACAC | 20 | Forward | 173 | RxLR |
| PcRx011_R2 |  |  |  |  |  | GGCAGTTTGGAGAAGTAACCTG | 22 | Reverse |  |  |
| PcRx012_F2 | -- | *PcAvh246/t* | -- | g24792 | g25726 | CTACTGGTGCTTTTGGTCATCG | 22 | Forward | 159 | RxLR |
| PcRx012_R2 |  |  |  |  |  | TTCTTCGCTGTCCATTTCGC | 20 | Reverse |  |  |
| PcRx013_F2 | *Pi Avr-blb2* | *PcAvh320* | g16030/g16031 | g27333 | g26879 | GTCACAGGTCCAGAAGATGCT | 21 | Forward | 208 | RxLR |
| PcRx013_R2 |  |  |  |  |  | CTTCGACAGCCTCAAACTCTG | 21 | Reverse |  |  |
| PcRx014_F2 | -- | *PcAvh136* | g16973/g16975 | g26851 | g26755 | CTGACGAGGAAGAAGACGAAGAAG | 24 | Forward | 181 | RxLR |
| PcRx014_R2 |  |  |  |  |  | GGGCTGTAGGGGTTGTAGTTAG | 22 | Reverse |  |  |
