## Supplementary material for "Comparative analysis of host-associated variation in *Phytophthora cactorum*": Supp. Table 6 Known Avrs.docx

| **KNOWN  AVR** | **GENBANK ACC.** | **AA (AFTER SP)** | **PcAvh ID** | **P414 GENE ID**  **(Pcac1_)** | **P414 EXP.** | **R36/14 GENE ID**  **(PC123_)** | **R36/14 EXP.** | **17-21 GENE ID**  **(PC128_)** | **17-21 EXP.** | **% SIMILARITY** | **NOTES** |
| --- | --- | --- | --- | --- | --- | --- | --- | --- | --- | --- | --- |
| PiAvr1 | DS028168.1 | 23-209 | PcAvh210/1/2 | g15762 | Yes | g15192 | -- | g15282 | -- | 52.5 |  |
| PiAvr2 | XM_002902939.1 | 21-117 | -- | -- | -- | -- | -- | -- | -- | -- |  |
| PiAvr3a | AEH27535.1 | 22-147 | PcAvh258 | -- | -- | g19522 | Yes | -- | -- | 57.9 |  |
| PiAvr3b | XM_002998411.1 | 20-254 | PcAvh377/78/79/80/81/82 | g11509/g11873/ g11938 | No | g22203/g15619/ g6721 | -- | g10109/g18031/ g4369 | -- |  |  |
| PiAvr4 | EF672355.1 | 25-287 | PcAvh356/7 | g4103 | No | g3490 | -- | g15741 | -- | 52.6 | No start codon in R36/14 & 17-21, genes called wrong |
| PiAvrblb1 | EEY61733.1 | 22-152 | PcAvh394 | g3106 | Yes | g26318 | -- | g18559 | -- | 41.7 | Also known as ipiO |
| PiAvrblb2 | XM_002895872.1 | 23-100 | PcAvh320 | g16030/1 | Yes | g27333 | No | g26879 | Yes | 26.4 |  |
| PiAvrvnt1 | XM_002997111.1 | 24-154 | PcAvh428/9 | g7005 | No | g16330 | -- | g11784 | -- | 56.8 |  |
| PiAvrSmira1 | KX887490.1 | 24-238 | PcAvh266/7/8 | g15770 | Yes | g18579 | -- | g6728 | -- | 35.6 |  |
| PiAvr8  (PiAvrSmira2) | XM_002904498.1 | 21-244 | -- | g3290 | No | g17704 | -- | g16289 | -- | 50.0 | Not called as RxLR in P414, introns in R36/14 & 17-21 |
| PiAvramr1 | XM_002904507.1 | 22-287 | PcAvh335 | g9245 | Yes | g12607 | Yes | g9824 | Yes | 61.3 |  |
| PiAvramr3 | XM_002895186.1 | 23-339 | PcAvh092 | g26927 | No | g24106 | No | g11927 | No | 62.1 |  |
